# *In situ* lipidomics of *Staphylococcus aureus* osteomyelitis using imaging mass spectrometry

**DOI:** 10.1101/2023.12.01.569690

**Authors:** Christopher J. Good, Casey E. Butrico, Madeline E. Colley, Katherine N. Gibson-Corley, James E. Cassat, Jeffrey M. Spraggins, Richard M. Caprioli

## Abstract

Osteomyelitis occurs when *Staphylococcus aureus* invades the bone microenvironment, resulting in a bone marrow abscess with a spatially defined architecture of cells and biomolecules. Imaging mass spectrometry and microscopy are invaluable tools that can be employed to interrogate the lipidome of *S. aureus*-infected murine femurs to reveal metabolic and signaling consequences of infection. Here, nearly 250 lipids were spatially mapped to healthy and infection-associated morphological features throughout the femur, establishing composition profiles for tissue types. Ether lipids and arachidonoyl lipids were significantly altered between cells and tissue structures in abscesses, suggesting their roles in abscess formation and inflammatory signaling. Sterols, triglycerides, bis(monoacylglycero)phosphates, and gangliosides possessed ring-like distributions throughout the abscess, indicating dysregulated lipid metabolism in a subpopulation of leukocytes that cannot be discerned with traditional microscopy. These data provide chemical insight into the signaling function and metabolism of cells in the fibrotic border of abscesses, likely characteristic of lipid-laden macrophages.

## INTRODUCTION

Infectious diseases can be debilitating and deadly when not treated properly. Even with advanced modern medicines and antibiotics, some infections carry poor treatment success rates due to antimicrobial resistance of the pathogen and tissue inflammation that can lead to insufficient drug delivery.^1^ Osteomyelitis, characterized by bone inflammation and most commonly caused by bacterial invasion, is one such disease with a 20% probability of treatment failure and high likelihood of recurrent infections.^1,2^ Although the incidence rate is estimated to be ∼22 per 100,000 person-years, comorbidities like diabetes increase infection susceptibility and underscore the clinical relevance of this disease in an increasingly complex patient population that frequently carries medical comorbitities.^3^

Bacteria gain access to the bone microenvironment by means of the bloodstream, a contiguous soft tissue infection, or a direct structural opening resulting from traumatic fractures or surgical complications.^4,5^ The majority of osteomyelitis cases are caused by *Staphylococcus aureus*, a Gram-positive opportunistic pathogen notorious for its antimicrobial resistance and ability to infect a variety of other mammalian organ systems, including but not limited to the skin, heart, lung, and kidney.^3,6,7^ Once exposed to the cellular-rich bone marrow within the intramedullary cavity of long bones, *S. aureus* employs a broad range of virulence factors to colonize, proliferate, and disseminate throughout the organ.^5,8^ After affixing to cells and extracellular matrix in the bone, *S. aureus* triggers the recruitment of innate leukocytes and deploys toxins and immunoevasive factors to protect itself from immune defenses. Over multiple days, the resulting host-pathogen interactions promote the formation of an inflammatory lesion or abscess. The hallmark architecture of an abscess consists of a central bacterial microcolony, known as a staphylococcal abscess community (SAC), surrounded by layers of necrotic and viable neutrophils and leukocytes.^8^ In addition, an outer fibrous layer develops, effectively sequestering the necrotic abscess from healthier surrounding tissue. The maturation of an abscess establishes a structural barrier between viable leukocyte defenses and the proliferating colony. Vascular impairment accompanies abscess development and is hypothesized to be responsible for poor antibiotic delivery and treatment failure.^1^ If endogenous leukocytes or exogenous antibiotics do not adequately eliminate the SAC, *S. aureus* can escape the abscess and disseminate to other regions within or outside the bone.

Lipids serve critical roles for innate immune defenses and bacterial immune evasion. For example, *S. aureus* employs a virulence factor, multiple peptide resistance factor (MprF), that chemically modifies the existing phosphatidylglycerol pool in its cell membrane by attaching a lysine to its phospholipid headgroup, effectively inverting its membrane polarization to help repel positively charged antimicrobial compounds.^9,10^ Without this lipid reaction, *S. aureus* survival in the presence of neutrophils decreases, and virulence becomes attenuated in murine infection models.^9^ From the host perspective, lipids influence cellular structure and energy production while also serving as substrates for various lipid-mediated inflammatory signaling pathways. Of particular interest, ether lipids are characterized by an ether-alkyl or ether-alkenyl bond at the *sn*-1 position of the glycerol backbone. This unique subset constitutes roughly 20% of the phospholipid pool in mammals and is more heavily produced in leukocytes found throughout the bone marrow.^11^ Although their function has not been entirely elucidated, ether lipids are implicated in maintaining membrane homeostasis and activating inflammatory signaling pathways.^12^ One signaling ether phosphatidylcholine, called platelet-activating factor, can be synthesized by leukocytes and functions in an autocrine or paracrine manner to stimulate host defense processes.^13^ Furthermore, inhibition of ether lipid synthesis results in neutropenia and disease, indicating their importance for leukocyte survival and function.^14^ Other characterized lipid-mediated signaling pathways involve eicosanoids and specialized pro-resolving mediators, which are both oxidized products of polyunsaturated fatty acids (PUFAs). These pro-inflammatory or anti-inflammatory molecules are typically derived from arachidonoyl phospholipids found within the membranes of cells that respond to infection.^15^ Following receptor-mediated stimulation, the arachidonic acid is lysed from the precursor lipid and subjected to oxidation by enzymes like cyclooxygenase and lipoxygenase, and the resulting products stimulate or resolve inflammatory reactions.^16^ In the scope of osteomyelitis pathology and abscess formation, the molecular and cellular interactions are inherently spatially defined. Omics data provides mechanistic insight into cell populations, but significant information regarding cell environment is lost without spatial preservation of a diseased structure like an abscess. Uncovering the complex biological processes that drive osteomyelitis pathology requires multimodal technologies that consolidate omics data with spatial cellular analysis *in situ*. Matrix-assisted laser desorption/ionization (MALDI) imaging mass spectrometry (IMS) paired with microscopy provides spatiomolecular information that incorporates label-free, multiplexed omics data with cellular resolution histological analysis.^17,18^ Briefly, molecules are desorbed and ionized by laser irradiation across the surface of a tissue section at discrete x-y coordinates (pixels), and the intensity of a selected *m/z* value at each pixel is displayed as an ion image. Ion images can then be correlated, registered, or computationally fused with compatible microscopy modalities to provide morphological context for molecular distributions.^19–22^

MALDI IMS has been successfully applied to bacterial infection models to understand the molecular interactions at the host-pathogen interface, including infections caused by common human pathogens like *S. aureus*,^23–29^ *A. baumannii*,^30,31^ *C. difficile*,^32,33^ *M. tuberculosis*,^34,35^ *S. enterica*,^36^ and *P. aeruginosa*.^37^ Investigation of the spatially defined host-*S. aureus* interface in bone will provide insight into the molecular dynamics that drive the devastating progression of the disease. Herein, we demonstrate an imaging workflow for interrogating the lipidome of a *S. aureus* osteomyelitis model given the important role of these molecules in innate immune responses and abscess patterning. First, lipid atlases were created to identify specific markers of tissue types throughout the infected femur. The intensities of lipids are then reported for different regions of bone marrow abscesses including the SAC, inner neutrophilic region, and outer fibrotic region. Finally, lipids are discovered to associate with specialized cells in the fibrotic border of abscesses.

## RESULTS

### Multimodal Molecular Imaging of a *S. aureus* Osteomyelitis Model

An imaging workflow incorporating microscopy and MALDI IMS was applied to an osteomyelitis murine model to understand the molecular dynamics fundamental to the progression of *S. aureus* infection in bone.^38,39^ For the majority of experimental analyses, femur section replicates were collected from three infected and three mock-infected mice. Non-destructive, unlabeled fluorescence microscopy was leveraged to provide morphological images of tissues within and surrounding the section prior to MALDI IMS (**Figure 1A**). The autofluorescence signal of the infected bone marrow distinguished abscess regions and was more intense than the mock-infected bone marrow (**Figure S1A**). The inoculated *S. aureus* constitutively expressed GFP to determine the precise location of SACs within the intramedullary cavity.

**Figure 1.**
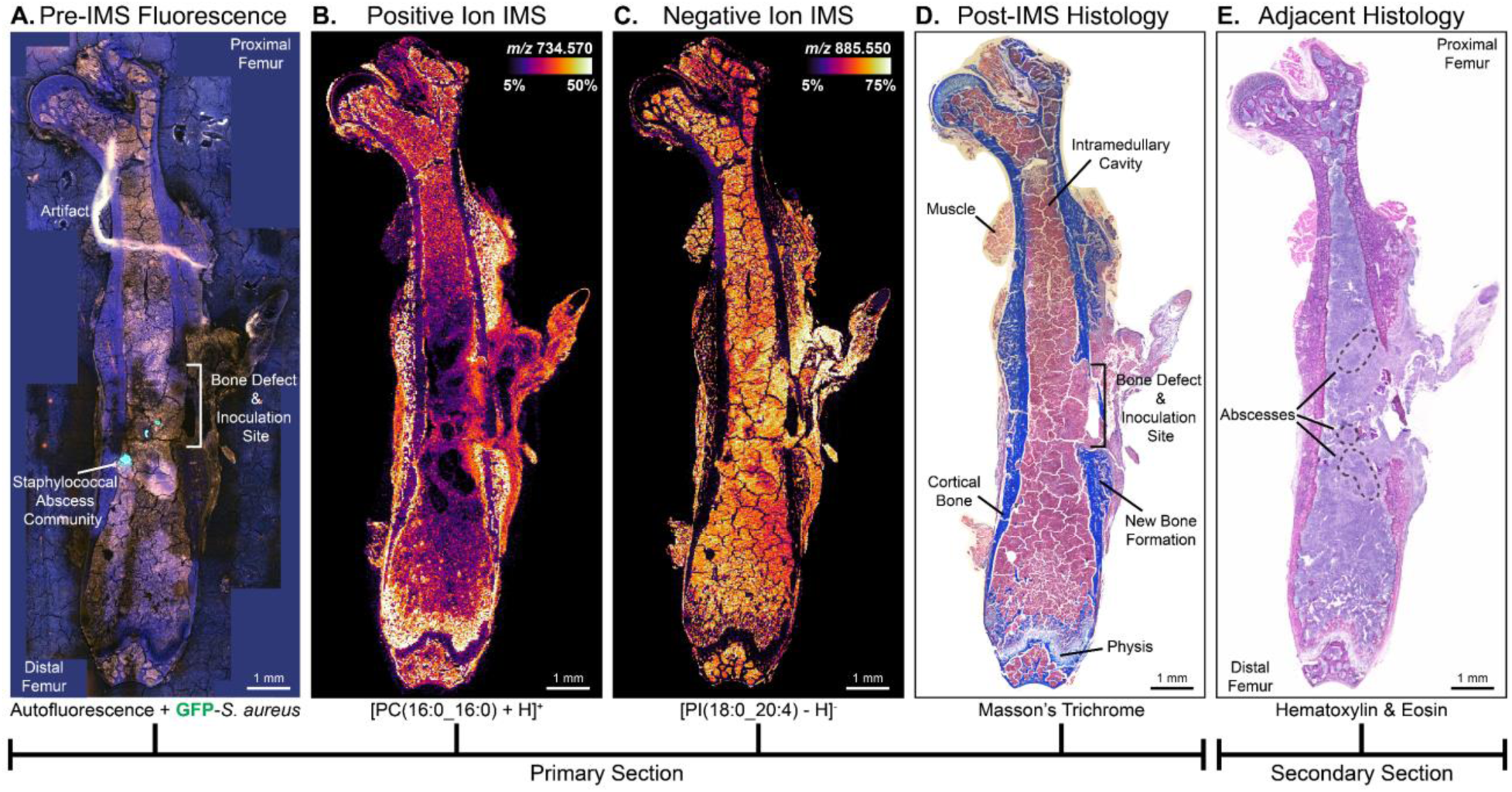
Microscopy and MALDI IMS are used to map and understand lipid distributions throughout a *S. aureus* infected femur. (A) Pre-IMS fluorescence microscopy captures autofluorescence signal from murine tissues and GFP fluorophore expression from *S. aureus*. (B,C) [PC(16:0_16:0) + H]^+^ and [PI(18:0_20:4) - H]^-^ are the two most abundant lipids detected in sequential MALDI IMS experiments. Data is acquired in positive and negative ion modes from the same section using a pitch offset strategy, yielding an effective spatial resolution of 20 μm. (D) Post-IMS histological staining with Masson’s Trichrome (MTC), or others, aids in tissue type identifications. (E) An adjacent section is fixed to prevent artificial bone marrow cracks and aids in cell type identifications throughout the intramedullary cavity. Annotations are provided where appropriate.

The MALDI matrix, 2,5-dihydroxyacetophenone (DHA), was selected to maximize lipid coverage by dual polarity ionization and applied on top of the tissue by sublimation to minimize both lipid delocalization and bone marrow damage. Two MALDI IMS experiments were performed sequentially on the same femur section by using a pitch offset strategy (**Figures 1B and 1C**). Briefly, positive ion data were acquired first to detect lipids such as phosphatidylcholines (PCs). The diameter of the laser burn marks was smaller than the stage pitch to leave unablated matrix between each ablation crater. The stage coordinates were then offset to enable a second laser ablation pass targeting previously unablated regions for negative ion analysis, yielding distributions of lipids such as phosphatidylinositols (PIs). [PC(16:0_16:0) + H]^+^ and [PI(18:0_20:4) - H]^-^ were the most abundant lipids detected throughout infected femur sections in positive and negative ion mode, respectively. This dual polarity approach expanded the coverage of identifiable lipids including those from the following additional classes: sphingomyelins (SMs), phosphatidic acids (PAs), phosphatidylethanolamines (PEs), phosphatidylserines (PSs), phosphatidylglycerols (PGs), cardiolipins (CLs), bis(monoacylglycero)phosphates (BMPs), sulfatides (SHexCers), gangliosides (GMs), triglycerides (TGs), and cholesterol esters (CEs), in addition to lysine (Lysyl-), lyso (L), and ether modifications (O-) of those lipids. Complete lists of identified murine and *S. aureus* lipids based on accurate mass measurements and liquid chromatography tandem mass spectrometry (LC-MS/MS) validation are reported (**Tables S1 and S2**).

Following MALDI IMS, staining using hematoxylin and eosin (H&E), Masson’s Trichrome (MTC), or Oil Red O (ORO) was performed on different section replicates of each femur to help characterize tissue pathology (**Figures 1D and S1B**). These histological stains emphasized cracking artifacts in the bone marrow that arise from tissue thawing steps prior to MALDI IMS.^38^ When preparing tissue sections adjacent to those analyzed by MALDI IMS, sections were fixed after removal from the cryostat and were not allowed to air dry throughout the staining protocols. This modified approach minimized bone marrow cracking so that some cell types could be better annotated (**Figure 1E**).

### Lipid Atlases of Tissue Types Throughout Infected Femurs

Efficiently characterizing the lipidome of tissue types throughout an infected femur began to offer molecular insight into bone infection processes. Unsupervised machine learning (e.g. *k*-means clustering) was employed to group pixels in an untargeted manner based on their spectral profile, and the resulting clustering data was used to perform segmentation where each color in a segmented image represents a cluster with a unique molecular profile. Segmented images for select tissue section replicates are shown for positive and negative ion analyses, where +*k* and -*k* reference positive and negative clusters, respectively (**Figures 2AI and 2AII**); all replicate images are also reported (**Figures S2AI and S2AII**). Segmented images serve as a representation of the molecular heterogeneity between clusters, which were primarily driven by different tissue types, cell types, and cellular processes throughout the femur section. For example, clusters +*k*5 and -*k*4 (red segments) were molecularly different than clusters +*k*9 and -*k*7 (gray segments); the former were associated with muscle, and the latter were associated with cortical bone, based on their correlation to histology. The average mass spectrum for a given cluster can be extracted to recapitulate the molecular heterogeneity that drives the clustering algorithm (**Figure S3**). There were apparent differences in segmentation patterns between positive and negative ion data due to the distinct molecular profiles generated by each polarity. The most abundant lipid in negative ion mode, [PI(18:0_20:4) - H]^-^, was ubiquitous throughout the femur and was suspected of driving the less-defined clustering. By removing this lipid from the analysis, negative ion clustering was more defined to tissue types, similar to positive ion data (**Figure S2B**).

Although healthier surrounding tissue types were still characterized by this atlas approach, bone- and infection-associated tissue types are the focus of this work. As seen from post-IMS histology, cluster segments corresponded well with annotated tissue types related to bone growth and wound healing in the distal and diaphysis regions of the femur **(Figures 2B and 2C**). The segments associated with cluster +*k*7 (yellow) and clusters -*k*5-6 (yellow) marked the chondrocyte-rich physis or growth plate of long bones. New bone was represented by cluster +*k*1 (purple) and developed in response to the surgical defect trauma.^40^ This tissue was found adjacent to the distal physis and adjacent to existing mineralized cortical bone near the defect-inoculation site. The periosteum, marked by cluster +*k*2 (orange), is a thin superficial layer covering the outer cortical bone. This segment was enlarged from the distal end to the defect-inoculation site, marking a fibrous connective tissue consistent with a hyperplastic response secondary to injury. Arterial endothelium and connective tissue near the joints like synovium were also marked by cluster +*k*2 (orange). These bone-associated clusters were present in mock-infected samples, but their segment areas and thus tissue sizes differed. Of note, new bone still formed in a mock-infected femur because a defect was created; however, the resulting bone damage appeared to be less severe because there was no infection to interfere with physiological bone healing.

**Figure 2.**
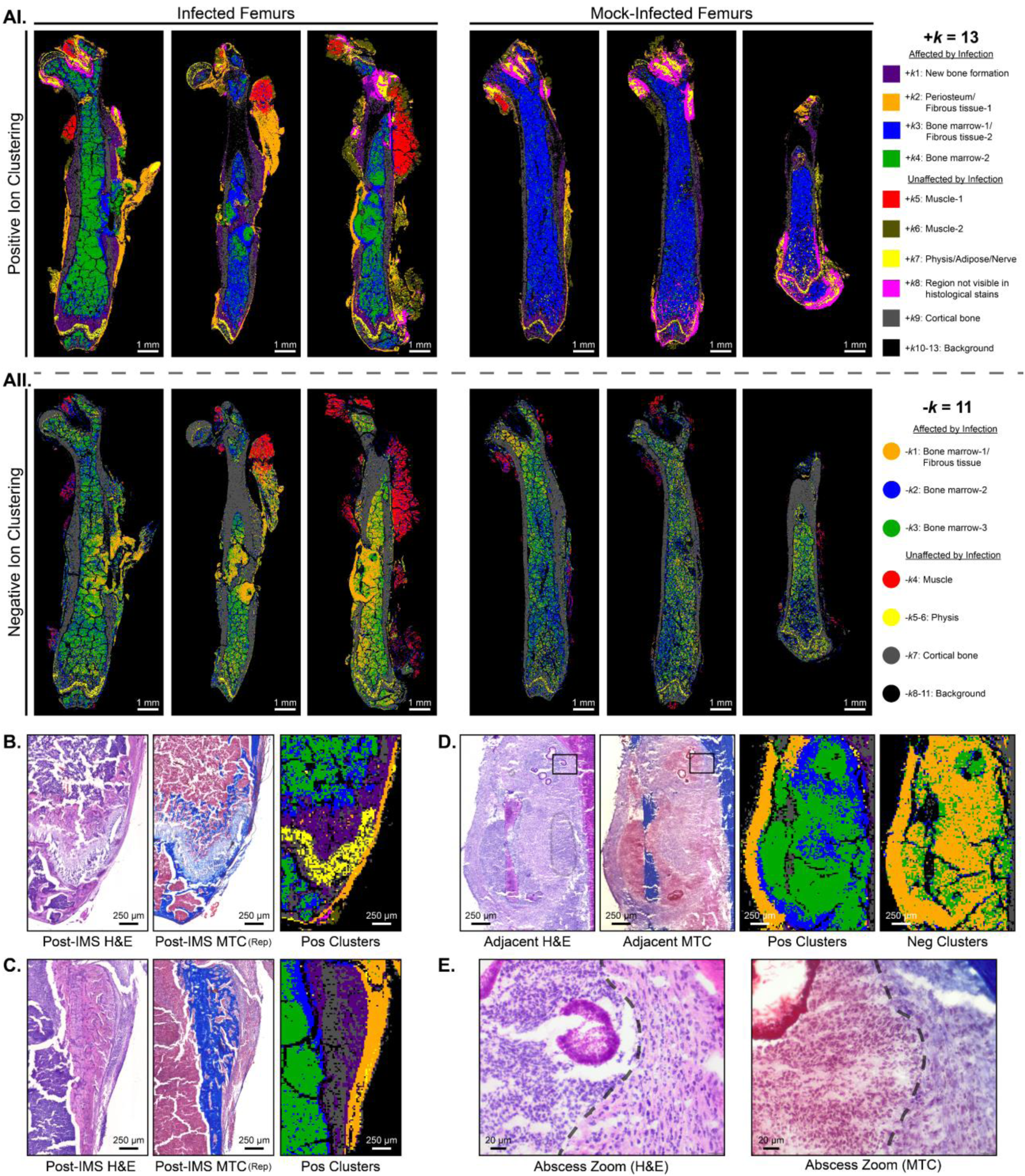
Machine learning and histology are leveraged to interrogate the murine lipidome. (A) *k*-Means clustering results and segmented images for separate (AI) positive and (AII) negative ion data reveal distinct pixel clusters that are associated with tissue types. A single section replicate from each infected and mock-infected femur are shown, and cluster labels on the right are supported by histology. (B,C) Positive ion clusters mark tissue types like the physis (yellow), new bone formation (purple), and periosteum/fibrous tissue (orange) in the (B) distal femur and (C) femoral diaphysis. Corresponding zooms of post-IMS histological stains are shown, one of which is a stain of a MALDI IMS section replicate (Rep). (D) Adjacent histology is used to help explain the molecular heterogeneity driving the bone marrow clusters in positive and negative ion modes. (E) Higher magnification of the H&E and MTC stains demonstrate the demarcation (dotted line) of degenerate or viable neutrophils to the left of the line, and fibrous tissue to the right of the line. The spatial resolution for all clustering datasets is 20 μm.

Cellular annotations from adjacent histology were leveraged to help define the intricacies of the cellular populations and tissue pathology present at the inoculation site and throughout the intramedullary cavity (**Figure 2D**). As previously stated, cluster +*k*2 (orange) associated with fibrous tissue outside of the intramedullary cavity, which was further supported by a thick collagenous matrix more clearly represented in adjacent H&E and MTC stains. Within the intramedullary cavity, the cellular composition of a spherical abscess can be visualized, and a dividing line was added to separate the two global regions of an abscess (**Figure 2E**). Cluster +*k*4 (green) largely associated with degenerate or viable neutrophils to the left of the dividing line or inner portion of the abscess. Cluster *+k*3 (blue) marked spindle cells, likely fibroblasts, and other bone marrow cells and leukocytes embedded in a collagenous matrix to the right of the dividing line or outer portion of the abscess. Although not consistent across all replicates, cluster -*k*3 (green) more closely associated with cluster +*k*4 (green) and its inner abscess neutrophil populations. Cluster -*k*1 (orange) appeared to be a general combination of cluster +*k*2 (orange) and cluster +*k*3 (blue), both of which marked fibrous tissue. This positive cluster deviation was driven by a sodium distribution outside the intramedullary cavity, which only impacted positive ion data. In both polarities, there were more green pixels in infected femurs compared to mock-infected femurs, suggesting a change in detectable lipids or abundance of those lipids. Based on histopathological assessment, this difference in lipid profile was most likely a result of the high influx of inflammatory cells discovered in abscesses and throughout the infected intramedullary cavity.

### Individual Lipid Markers of Tissue Types

To distinguish lipids that are primarily distributed to single clusters and thus candidate markers of the different tissue types present in the femur, the ratio of the mean intensity within a cluster versus the mean intensity across the whole tissue section was analyzed for each ion. Some of the most specific lipids with the highest ratio for each cluster are displayed as overlaid ion images (**Figures 3A and 3B**). A complete list of lipid markers for each cluster or identified tissue type is provided and organized by their polarity and association with osteomyelitis pathology (**Tables S3-S6**). Focusing on the impact of bacterial invasion into the bone marrow, trends developed in the top lipids identified in the bone marrow clusters. Cluster +*k*4 (green) and cluster -*k*3 (green) contained a plethora of ether lipids and sphingolipids while cluster +*k*3 (blue) and cluster -*k*1 (orange) consisted of many glycerophospholipids containing PUFAs. Changes to the abundances of these lipids in different cellular regions of abscesses were interrogated later.

**Figure 3.**
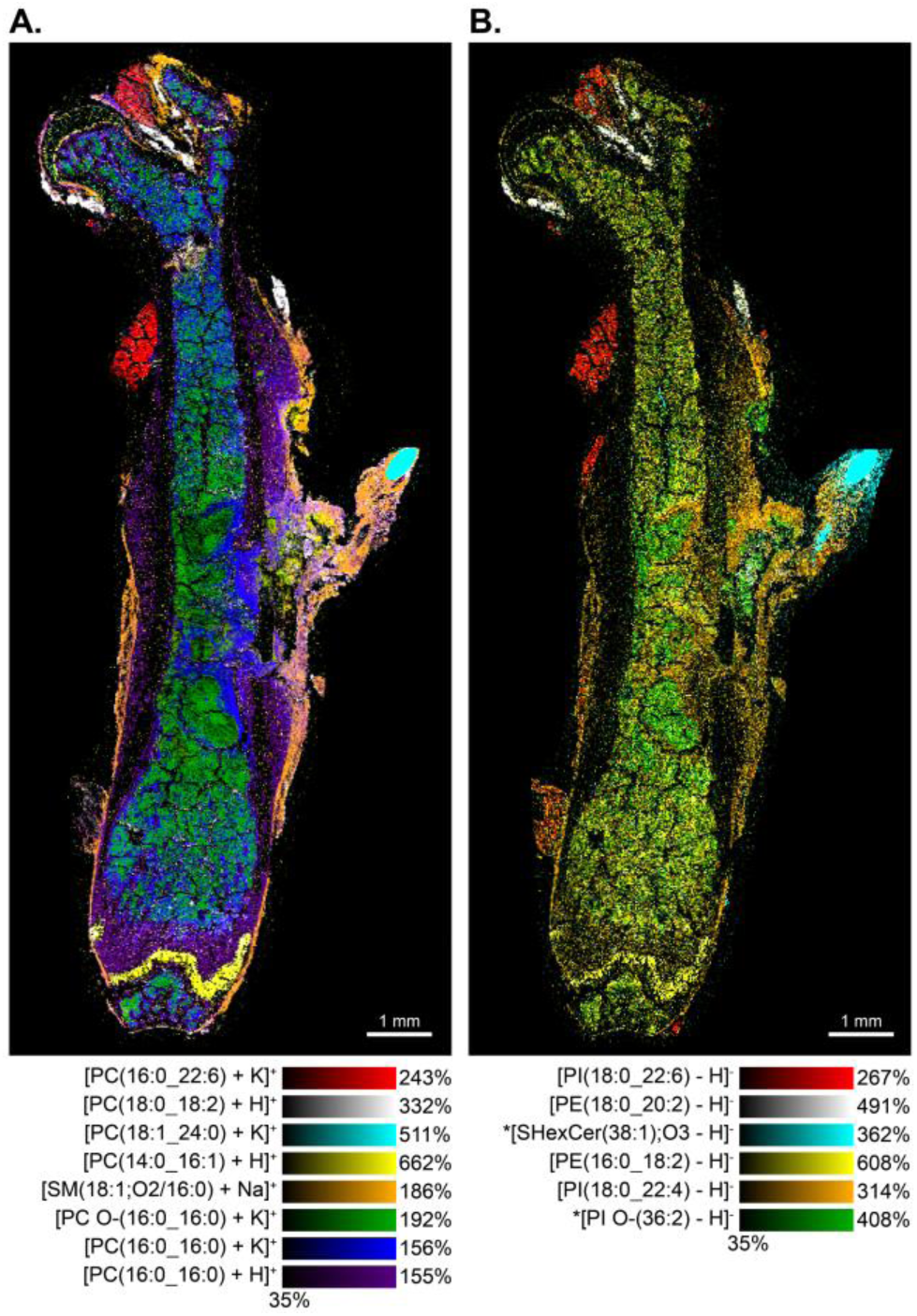
Lipids specific to tissue types are discovered from the *k*-means clustering analysis. (A) An overlaid image of eight positive ions displays discrete distributions to soft tissues surrounding and within the infected femur. The ions’ *m/z* (Top to Bottom) are 844.526, 786.601, 910.667, 704.524, 725.558, 758.548, 772.526, and 734.570. (B) An overlaid image of six negative ions highlight similar distributions. The ions’ *m/z* (Top to Bottom) are 909.550, 770.569, 850.572, 714.508, 913.580, and 847.570. The spatial resolution for the datasets is 20 μm, and hotspot removal intensity scaling was employed due to the wide range of ion intensities. * in ion image labels denotes preliminary identifications.

In positive ion mode, the periosteum and fibrous tissue outside of the intramedullary cavity possessed a high abundance of sodiated lipids, and the bone marrow was mainly dominated by potassiated lipids due to the distributions of salt throughout the femur. Even though salt washing induces additional morphological bone marrow damage, an experiment was conducted to remove endogenous salt leading to preferential formation of protonated lipids. As expected, the ion images from washed samples appeared simplified with increased intensity for protonated species and loss of sodiated and potassiated adducts (**Figure S4**). Clustering was performed on the washed sample, and even without adduct competition, there was enough lipid diversity to drive similar clustering results compared to unwashed tissue (**Figure S2C**). Since the primary focus of this study is bone marrow infection, potassiated lipids in the bone marrow were analyzed moving forward.

### Targeted Investigation of the Lipidome of Staphylococcal Abscess Communities

Once the murine lipidome was extensively mapped throughout the femur, more targeted methods were selected to interrogate the immediate host-pathogen interface and individual abscesses in the bone marrow. Five SACs were present near the bone defect and inoculation site of a chosen sample as seen from the fluorescence microscopy image (**Figure 4A**). After registering the two imaging modalities, regions of interest were drawn around the expressed GFP signal, in green, and around the whole inoculation site, in red (**Figure 4B**). Average mass spectra derived from the pixels within the green and red regions were overlaid to highlight several high intensity peaks that were unique to the SACs (**Figure 4C**). The most abundant bacterial lipid was [PG(32:0) - H]^-^, and its ion image confirmed its specificity to the SACs. Since prokaryotes are known to synthesize and utilize odd-chain fatty acids, it was unsurprising that [PG(31:0) - H]^-^ was also detected in the SACs. Lysine modifications of PGs were also discovered in the SACs, albeit with lower intensity in the higher *m/z* range (*m/z* 820-865, **Figure 4D**). The detection of Lysyl-PGs indicates that the membrane lipid conversion by MprF is conserved in this bone infection model.^28^ The main phospholipid constituents of *S. aureus* membranes are PGs, Lysyl-PGs, and CLs.^41^ Lipids belonging to all these classes were preliminarily identified in this study and were consistent with other studies that have used mass spectrometry (**Table S7**).^28,42,43^

**Figure 4.**
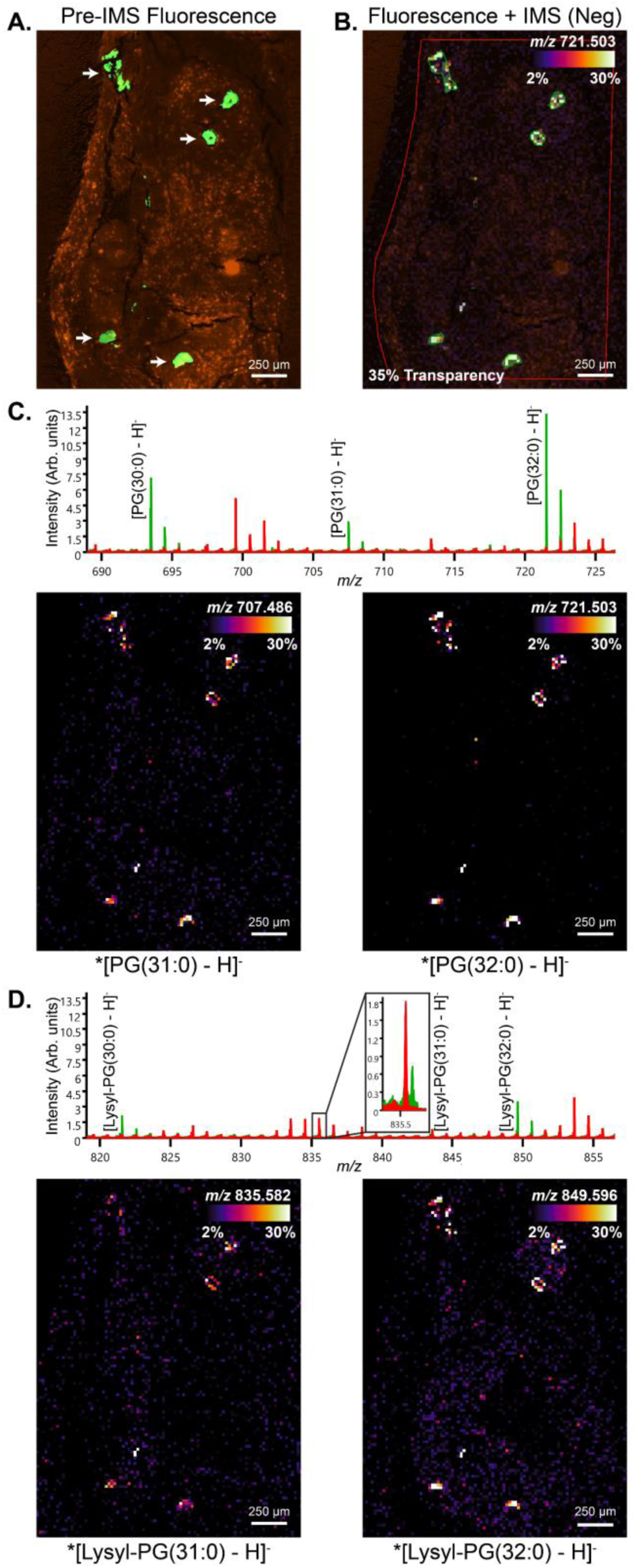
Imaging workflow enables targeted investigation of *S. aureus* lipids. (A) Fluorescence microscopy highlights five established SACs (white arrows) that are expressing GFP in the autofluorescent bone marrow. Intensity scaling for channels is adjusted. B) Microscopy is registered with 20 μm MALDI IMS data and regions of interest are more accurately drawn around SACs (green) and surrounding bone marrow (red). An ion distribution associated with *S. aureus* overlays the microscopy with 35% transparency to demonstrate registration accuracy. (C) Overlaid average mass spectra from the green and red extracted pixels highlight the most abundant PGs in the SACs. Peak annotations are provided where appropriate, and representative ion images recapitulate their specificity to bacteria. (D) Spectra and ion images are also shown for another class of *S. aureus* lipids, lysyl-PGs. * in ion image labels denotes preliminary identifications.

### Changes in Lipid Abundance in Individual Bone Marrow Abscesses

The goal of this analysis was to understand how lipids distributed throughout cellular regions of individual abscesses. An abscess of similar area was selected for each femur, and measurements of the same abscess from section replicates contributed to the replicate average (**Figure 5A**). The pre-IMS fluorescence image and post-IMS H&E stain showed distinct morphological changes associated with the outer and inner portions of the abscess, recapitulating the cell populations described previously (**Figures 5B and 2E**). A region of interest was drawn to encapsulate both abscess structures. Lipids abundant in the outer portion (e.g. [PC(38:4) + K]^+^) and inner portion (e.g. [PC O-(34:1) + K]^+^), in addition to ions corresponding to bacteria and background cracks, were all used to cluster regions of the abscess (**Figure 5C**). The same methods were repeated for negative ion data using [PE(38:4) - H]^-^ and [PI(36:3) - H]^-^, and clustering results were similar to the positive ion analysis (**Figure 5D**). All clustering replicates, and the location of the abscesses within the intramedullary cavities are also provided (**Figures S4A-C**). Bar graphs were used to compare the mean intensity of lipids between the outer and inner portions of abscesses (**Figures 5E and 5F**). Lipids that were significantly more abundant in the inner portion were SMs, ether PCs, ether PAs, PIs, PSs, CerPs, ether PIs, and ether PEs. There were additional lipids that followed similar trends but were not biologically significant due to abscess and replicate variation (**Figure S4D and S4E**). The lipids that were significantly more abundant in the outer portion of abscesses were PCs, PAs, LPCs, SMs, PIs, PEs, and ether PEs; the majority of which possessed high degrees of unsaturation. The two lipids that were chosen to cluster the outer portion, [PC(18:0_20:4) + K]^+^ and [PE(18:0_20:4) - H]^-^, contained the PUFA, arachidonic acid [FA(20:4)]. Leveraging LC-MS/MS experiments, 9 out of the 23 significant lipids with this abscess distribution contained arachidonic acid. Other PUFAs like docosatetraenoic [FA(22:4)], docosapentaenoic acid [FA(22:5)], and docosahexaenoic acid [FA(22:6)] were also detected in MS/MS spectra for many of these significant lipids.

**Figure 5.**
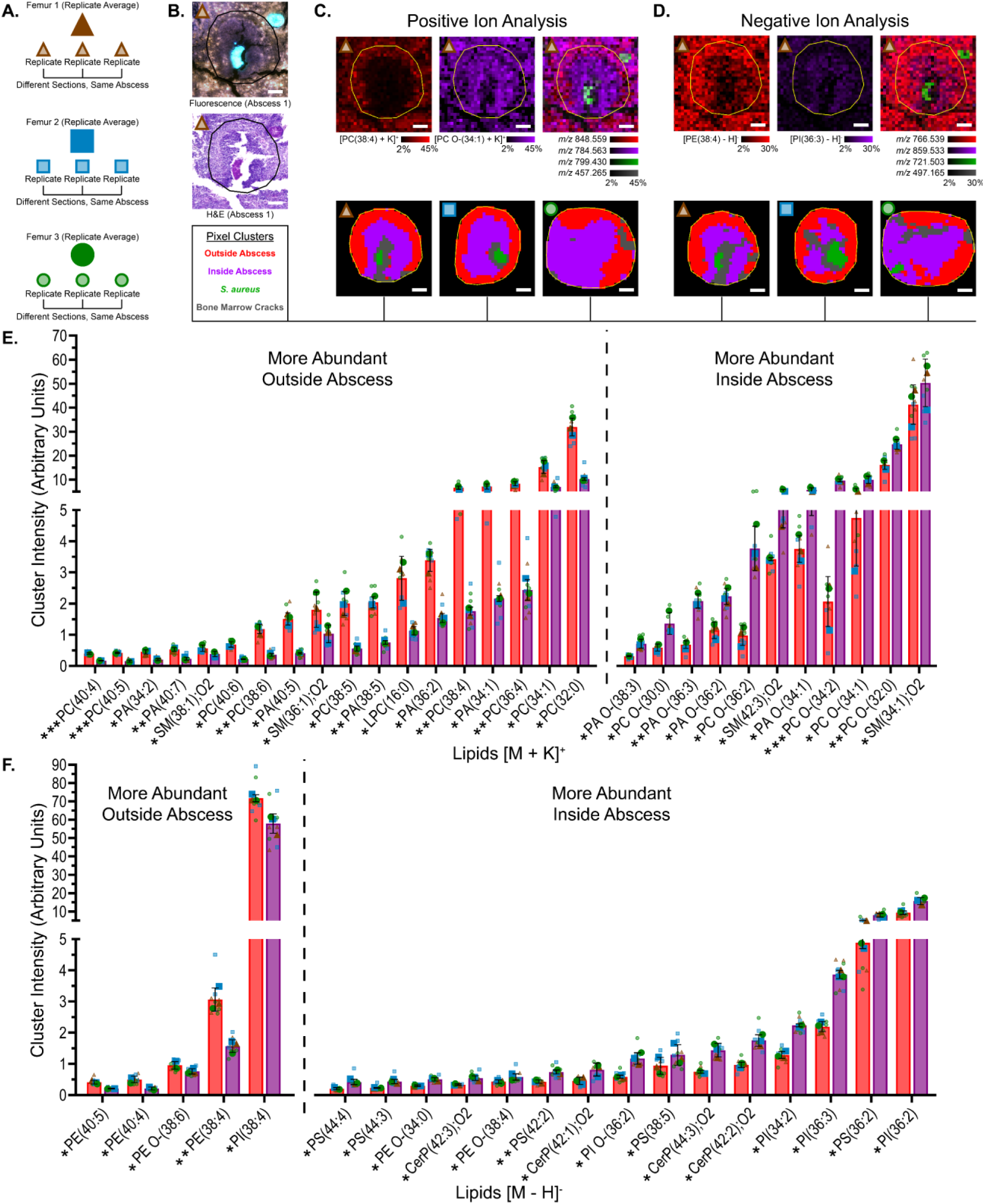
Clustering-assisted analysis of single abscesses reveals differences in lipid abundances between the outer and inner portions of bone marrow abscesses. (A) Schematic of experimental design shows single abscesses chosen from each femur, which is represented by different shapes. An outlined shape corresponds to a section replicate measurement of each abscess, and a solid shape corresponds to the average of section replicates for the abscess. (B) A pre-IMS fluorescence image and post-IMS H&E stain highlight visual changes between the two abscess regions contained within the drawn region of interest. (C,D) For (C) positive and (D) negative ion analysis each, four ion distributions (Top) contributes to the clustering algorithms, shown as individual and overlaid 20 μm ion images, and clustering results (Bottom) for each femur are reported. Scale bars represent 100 μm. (E,F) Bar graphs show the statistically significant differences in lipid abundances between the outside abscess pixels (red cluster) and inside abscess pixels (purple cluster) for each (E) positive and (F) negative ion analysis. The mean, with biological standard deviation, for three femurs is displayed as a bar with colors corresponding to the derived cluster. Section replicate measurements and replicate averages (one femur) are displayed on the graph. A paired T-test is used to test statistical differences, with *<0.05, **<0.01, ***<0.005 denoted below the graph on the x-axis labels.

Although this analysis targeted single abscesses, it was also beneficial to understand the abundances of these lipids in the total infected bone marrow compared to the mock-infected bone marrow. Most lipids that were more abundant inside the abscess were more abundant throughout the entire intramedullary cavity compared to its mock counterpart (**Figure S6**). These lipids were associated with more than neutrophils on the inside of established abscesses; they were detected in bone marrow cells and inflammatory cells found throughout the infected intramedullary cavity, which potentially indicates a lipidome shift in response to infection-induced myelopoiesis. On the contrary, some lipids that were more abundant outside the abscess were less abundant compared to the mock, such as [PC(16:0_16:0) + K]^+^ and [PI(16:0_18:1) - H]^-^. These lipids could either be substrates for conversion products or membrane components of bone marrow cells that were not as populated in infected femurs.

### Spatial Heterogeneity of Bone Marrow Lipids

Clustering entire femurs or clustering single abscesses did not capture the subtle heterogeneity in lipid distributions throughout the infected bone marrow. To account for these shortcomings, principal component analysis (PCA) was performed on pixels associated with the bone marrow from all infected sections (**Figure 6**). For positive ion mode, the loadings plot showed a clear separation among identified lipids for Component 1, which correlated well with findings from previous clustering analyses. (**Figure 6AI**). Lipids that are largely outside the abscesses consisted of a positive value for Component 1, and lipids inside the abscesses possessed a negative value. Component 3 differentiated lipids at the intersection between the inner neutrophilic and outer fibrotic abscess regions, termed here as the abscess border. A negative value Component 1 and positive value Component 3 defined a unique population of lipids with a distinct ring-like distribution, as seen from the most differentiated lipid, [PC O-(16:0_20:4) + K]^+^ (**Figure 6AII**). A positive value Component 1 and positive value Component 3 separated lipids, such as [PC(18:0_20:4) + K]^+^, that were largely abundant outside the abscesses but still maintained an intense concentration near the abscess border.

**Figure 6.**
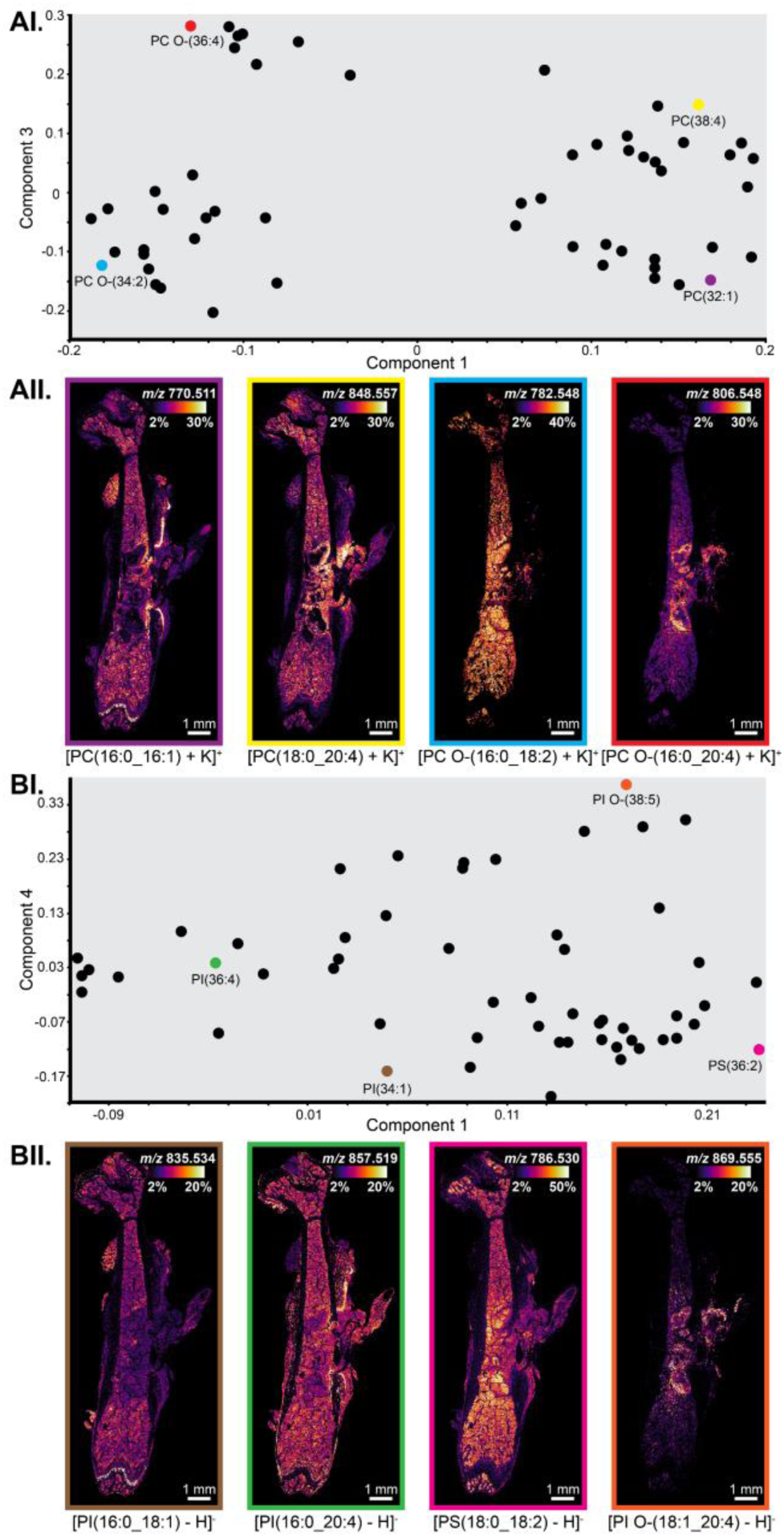
Principal component analysis (PCA) emphasizes the spatial heterogeneity of lipids detected throughout the intramedullary cavity. (A,B) Pixels associated with bone marrow in all nine infected samples are used in the analysis. (AI) Positive ion lipids (+K) separate by Component 1 and Component 3 as shown in the loadings plot. (AII) Ion images from the annotated and colored loadings exemplify the variation in distributions explained by these components. (BI) Negative ion lipids (-H) separate by Component 1 and Component 4 as shown in the loadings plot, and (BII) select ion images are also displayed. Each Component 1 is shown to highlight the largest variation and Components 3 and 4 are selected to represent the ring-like distribution. The spatial resolution for the datasets is 20 μm.

Differences observed in PCA of negative ion data were much less pronounced than the positive ion results. Still, Component 1 separated lipids based on their propensity to be detected in the inner or outer portion of an abscess, while Component 4 differentiated lipids with a similar ring-like distribution at the border (**Figure 6BI**). [PI O-(18:1_20:4) - H]^-^ and [PS(18:0_18:2) - H]^-^ represent lipid distributions that were abundant inside the abscess, with or without the pronounced ring (**Figure 6BII**). The cell types and cellular processes driving heterogeneity in lipid distributions, for example [PI(16:0_18:1) - H]^-^ and [PI(16:0_20:4) - H]^-^, are not fully understood, but the fibrous tissue within and outside the intramedullary cavity appear to be involved. The unique ring-like distributions and cell populations of the abscess border were further examined.

### Investigation of Lipids with Ring-like Distributions at the Abscess Border

Positive and negative ion data were acquired on adjacent sections at higher spatial resolution (10 µm) to target cell populations throughout a large abscess within the intramedullary cavity. To determine other lipids that localize to the ring-like distributions of [PC O-(16:0_20:4) + K]^+^ and [PI O-(18:1_20:4) - H]^-^, Pearson’s correlation coefficients (PCCs) were calculated between these lipids and other lipids detected in positive and negative ion modes, respectively (**Figures 7AI and 7AII**). Lipids with higher PCCs were more likely to have a similar pronounced ring-like distribution; however, high PCCs did not always equate to a pronounced ring-like distribution since the starting two lipids were also present in other regions of the abscess. For example, [SM(42:2);O2 + K]^+^ had a high 0.544 PCC, and it was abundant throughout the abscess, including the border, but it did not have a pronounced ring-like distribution. Lipid distributions that were pronounced and highly specific to the border were labeled with a solid bullet point for clarity (**Figures 7AI and 7AII**). Conversely, [PE O-(18:1_18:1) – H]^-^ had a lower 0.131 PCC and is an example of a lipid that was absent from the fibrous tissue (**Figure S6BII**). The most correlated ion images in both polarities were arachidonoyl ether glycerophospholipids (**Figure 7B**). Other highly correlated lipid classes included lyso ether lipids, lyso glycerophospholipids, TGs, CEs, BMPs, and GMs (**Figure 7C**). Based on PCCs and visual inspection of the high spatial resolution ion images, it was clear that these lipid classes were not necessarily associated with the same cell populations, but they were all closely associated with the general abscess border. Trapped ion mobility spectrometry (TIMS) was leveraged in an imaging experiment to confirm the identification of BMPs, which are unique isomers of PGs where the *sn*-2 fatty acid of the glycerol backbone is located on the glycerol headgroup (**Figure S7**). All of these lipid distributions helped inform the lipidome of cells at the abscess border.

**Figure 7.**
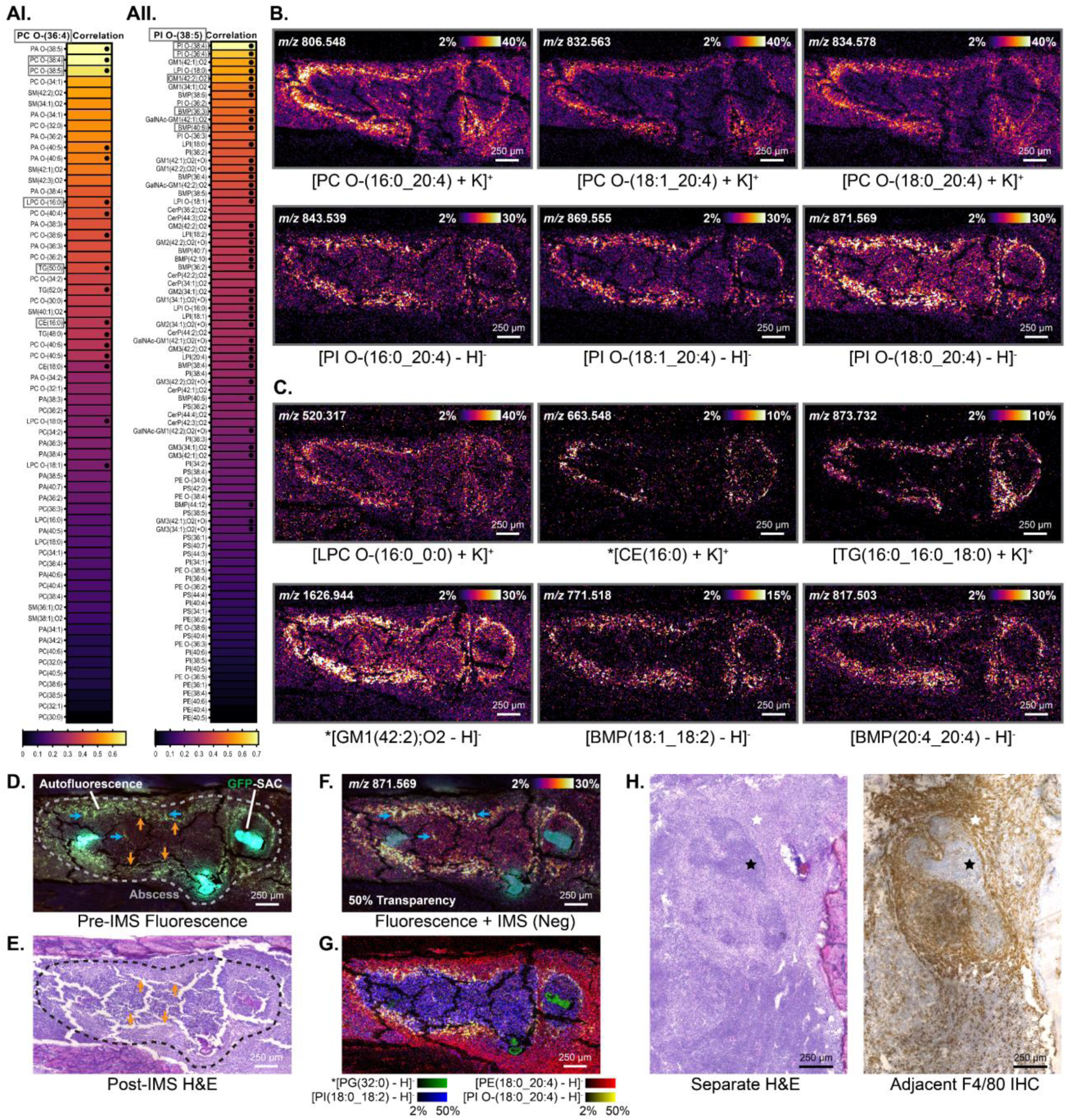
Unique lipid classes possess similar ring-like distributions associated with cells at the abscess border. (A) Pearson correlation tests highlight lipids (+K or -H) that correlate with the distribution of (AI) [PC O-(16:0_20:4) + K]^+^ and (AII) [PI O-(18:1_20:4) - H]^-^ within and surrounding a multi-SAC containing abscess (labeled in D & E). Black dots on the linear coefficient scale are supplemented to emphasize lipids with pronounced ring-like distributions as seen from their ion images. (B) Arachidonoyl ether glycerophospholipids are the most correlated lipids in both polarities. (C) Additional lipid classes like lyso lipids (L-), cholesterol esters (CEs), triglycerides (TGs), monosialo-gangliosides (GM1s) and bis(monoacylglycero)phosphates (BMPs) are also highly correlated. The spatial resolution for the datasets is 10 μm. (D) A pre-IMS fluorescence image shows autofluorescence signal (blue arrows mark hotspots), mostly in the GFP channel, derived from the abscess border morphology. (E) A post-IMS H&E stain with cracks (orange arrows) helps orient this autofluorescent signal mostly to the outer fibrotic region of the abscess. (F) A 50% transparent ion image of [PI O-(18:0_20:4) - H]^-^ is overlaid onto the fluorescence image to show their close association. (G) An overlaid ion image represents a more complete view of the lipid architecture of a bone marrow abscess. The ions’ *m/z* in reading order are 721.503, 766.539, 861.550, and 871.569. * in ion image labels denotes preliminary identifications. (H) H&E (Left) and F4/80 IHC (Right) staining is performed on a separate femur to confirm the presence of macrophages in the outer fibrotic region (white stars) surrounding the inner neutrophilic region (black stars).

To better understand the cellular composition of an abscess, genetically modified Catchup mice were used in this experiment where viable neutrophils were expected to express a tdTomato fluorophore.^44^ The autofluorescence signal of the bone marrow complicated the detection of the fluorophore, and identification of these reporter neutrophils could not be successfully validated. Nevertheless, the morphology of the abscess border was autofluorescing in the GFP channel, and a few hotspots were labeled with blue arrows in the pre-IMS fluorescence image (**Figure 7D**). Although not all, the majority of this signal was present in the outer fibrotic region of the abscess, as oriented by bone marrow cracks labeled with orange arrows in both the pre-IMS fluorescence image and post-IMS H&E stain (**Figures 7D and 7E**). The semi-transparent ion image of [PI O-(18:0_20:4) - H]^-^ was overlaid onto the fluorescence image to highlight how the ring-like distributions corresponded well to the autofluorescent border (**Figure 7F**). As seen from the overlaid ion images, the ring-like distribution (yellow) overlapped with lipids that marked the inner (blue) and outer (red) portions of an abscess, reestablishing the idea of a border region (**Figure 7G**). This observation was further supported by the fact that these border lipids did not significantly change between the two regions of the abscess (**Figures S5D and S5E**). Histopathology from literature and this study confirmed the presence of neutrophils and spindle cells near the border of a bone marrow abscess, and these cells could be contributing to the autofluorescence signal and lipid distributions.^45^ To better understand the contribution of macrophages, F4/80 IHC staining was performed adjacent to H&E staining of a separate femur (**Figure 7H**). The majority of this macrophage antigen was detected in the fibrous tissue and corresponded well with the previous 20 µm ring-like distributions (**Figures 6AII and 6BII**). Altogether, these unique lipids were strongly correlated to one another and were associated with the abscess border, where macrophages were detected and were most likely responsible for the ring-like distributions.

## DISCUSSION

Similar to many bone diseases and infectious diseases, osteomyelitis pathology is inherently spatially defined. Techniques that require sample homogenization will forfeit spatial information that is foundational for understanding disease development and progression. Spatial omic technologies like MALDI IMS can preserve the molecular and spatial architecture of disease features, but its application for bone diseases have been limited due to the practical difficulties of handling fresh-frozen bone.^38,46^ Our imaging workflow incorporating both MALDI IMS and microscopy yields unbiased and highly multiplexed lipidomics data from *S. aureus*-infected femurs in a spatially resolved and efficient manner. At some sacrifice to spatial resolution, the dual polarity data acquisition strategy from a single section is beneficial to reduce the sample quantity and thus lower sample preparation variability while maintaining optimal signal and maximum lipid coverage.

This study establishes lipid composition profiles for various tissue types related to the musculoskeletal system. Building lipid atlases provides chemical information that infers cellular function, and data can aid future investigations of tissue-specific pathologies. More importantly, our work provides critical insight into how lipids change in different cellular regions of a bone marrow abscess. Due to the high abundance of inflammatory cells identified via histology in the infected bone marrow, the significant changes in multiple ether lipids between inner and outer portions of abscesses and between infected and mock-infected femurs are believed to be associated with an influx of neutrophils and other inflammatory cells. Ether lipids are generally known to be an abundant component of leukocyte membranes.^11^ Furthermore, specific ether lipids have been identified in neutrophils and other leukocytes *in vitro* and have been spatially detected at sites of inflammation in animal models of bacterial infections.^28,47–49^ In our data, ether lipids have a high propensity to localize to bone marrow regardless of infection status, indicating some specificity to the hematopoietic stem cell niche under homeostatic conditions. Following bacterial invasion into the intramedullary cavity, myelopoiesis occurs in the same region, giving rise to a dense population of myeloid lineage cells.^50^ The associated increase in ether lipid abundances in the stem cell niche may indicate their involvement in myelopoiesis, either for membrane remodeling or lipid-mediated signaling. Due to the chemical and physical properties of ether lipids, we hypothesize that ether lipids are implicated in altering membrane fluidity for phagocytic function. More generally, these ether lipids could be defining lipid markers of a larger subset of myeloid lineage cells, not just neutrophils in the abscess.

Sphingolipid and acyl-glycerophospholipid classes are also abundant in the inner portion of abscesses and throughout the intramedullary cavity. The high abundance of specific SMs in the abscess-localized neutrophil population could indicate sphingomyelin synthase and ceramide kinase activities, which have both been shown to be crucial for neutrophil phagocytic activity.^51,52^ SMs have also been linked to direct activation of the transcription regulator NF-kB in leukocytes.^53^ Additionally, subpopulations of neutrophils in the abscess are hypothesized to undergo apoptosis. PS is synthesized and incorporated on the outer leaflet of apoptotic cell membranes for efferocytosis, so a high abundance of PS in these cells, such as [PS(18:0_18:2) - H]^-^, may allude to an apoptotic phenotype.^54^ Unlike previously published infection models, the bone-pathogen interface is at the site of myelopoiesis, yielding a unique molecular perspective into leukocyte development, recruitment, and function.

Our results also highlight a significant trend where lipids containing arachidonic acid and other PUFAs are less abundant in the inner neutrophilic regions compared to the outer fibrotic regions of abscesses. This trend could simply be a consequence of cell death that occurs in the inner portion of an established abscess. Arachidonoyl and other PUFA-containing lipids are also known to be a major source of the secondary fatty acid mediators that are involved in inflammatory signaling pathways, so a decrease in these precursor molecules in the abscess could rather indicate the liberation of the product fatty acid at an earlier timepoint of infection.^55,56^ Scott et al.^57^ used MALDI IMS to reveal the degradation of arachidonoyl PIs and PEs over the course of a bacterial infection in a murine spleen, and their work supports our findings and hypothesis that arachidonoyl lipids are degraded within abscess-localized cells to stimulate or resolve inflammatory processes. The downstream signaling consequences and conversion products of arachidonic acid are complex and not fully characterized in the context of abscess formation in osteomyelitis. For example, a single eicosanoid called prostaglandin E_2_ (PGE_2_) is converted from arachidonic acid by cyclooxygenases. During early inflammation and abscess development when neutrophils are recruited to bacteria stimuli, PGE_2_ is known to recruit and activate additional leukocytes.^58^ Miek et al.^59^ showed a significant increase of PGE_2_ in bone after an acute infection compared to a chronic infection, suggesting its early pro-inflammatory function. Still, PGE_2_ remains in a chronic setting and is known to also have a contrasting anti-inflammatory function by regulating cytokine production.^58,60^ Regardless of other eicosanoids, even the function of PGE_2_ is not entirely understood in osteomyelitis, especially when investigating localized cellular neighborhoods and not bulk organ expression. In all, lipid distributions presented here do not reveal the eicosanoids present or their pro-inflammatory or anti-inflammatory function, but they may indicate the specific arachidonic acid- and PUFA-containing substrates involved in lipid-mediated inflammatory signaling pathways that occur prior to the 14 dpi timepoint. Earlier timepoints should be investigated further to understand the development of inflammatory responses in the bone marrow.

Ether lipids that contain arachidonic acid distinguish a unique population of cells and leukocytes that reside within an abscess region termed here as the abscess border. We have shown evidence of collagen, spindle cells, neutrophils, and macrophages in this region, and we hypothesize that macrophages in the fibrous tissue are majorly responsible for the ring-like lipid distributions. Importantly, arachidonoyl ether lipids have already been detected in both neutrophil and macrophage cultures.^48,49,61^ A large quantity of arachidonic acid in the form of an ether glycerophospholipid at this timepoint potentially indicates that leukocytes are primed for future arachidonic acid cleavage for downstream signaling functions. The evidence of highly correlated lyso products, such as [LPC O-(16:0_0:0) + K]^+^, would suggest some phospholipase A2-mediated cleavage has already occurred in these viable cell populations.

Our results also reveal additional lipid classes with pronounced ring-like distributions at the abscess border, like neutral fat lipids, BMPs, and GMs. Given this reported lipidome, we believe macrophages with a lipid-laden phenotype, also known as foam cells, are present at the abscess border. Foam cell development is a critical event in disorders like Tuberculosis and atherosclerosis, where lipids like TGs and CEs accumulate intracellularly in the form of lipid droplets as a result of imbalances in neutral lipid uptake, synthesis, degradation, and efflux mechanisms.^62,63^ Foam cells develop in the presence of a pathogen or oxidized low-density lipoproteins (ox-LDL), like in granulomas or atheromas, respectively.^62^ Studies have reported a high abundance of TGs and CEs in foam cells in the context of other infection models.^64,65^ Additionally, Nicolaou et al.^66^ observed lipid droplet formation in macrophages following exposure to *S. aureus.* The phenotype is dependent on toll-like receptor 2 (TLR2) signaling and is similar to the more characterized mechanisms in Tuberculosis.^67^ This suggests the accumulation of lipid droplets in cells at the abscess border could be a direct result of recognizing *S. aureus*. Necrotic and inflammatory cells release damage-associated molecular patterns (DAMPs) that can activate TLR2 signaling, so crosstalk between the inner and outer portions of the abscess could also be responsible.^68^ Additionally, following receptor-mediated uptake of ox-LDL that is found in chronic inflammatory settings, lysosomes can become overloaded resulting in neutral lipid accumulation throughout the cell.^69^ In sum, the detection of TGs and CEs in the abscess border suggest the presence of intracellular lipid droplets, but the exact mechanism of their development is not confirmed in this osteomyelitis model.

The BMPs identified here correspond well with previous reports of BMPs possessing high degrees of unsaturation, including dual attachment of PUFAs such as [BMP(20:4_20:4) - H]^-^.^70^ BMPs have been implicated in various lysosomal storage disorders, supporting the hypothesis that cells at the abscess border have overloaded lysosomes.^71^ Although BMPs are not completely understood to be a cause or consequence of foam cell development, they are closely associated with dysregulation of cholesterol homeostasis and the formation of the lipid-laden phenotype.^72^ Glycosphingolipids, like gangliosides, are present in foam cells of atherosclerotic tissue, and accumulation within the cell increases LDL uptake and directly affects cholesterol homeostasis.^73–75^ Additionally, gangliosides and BMPs have been linked in models of gangliosidoses, which are neurodegenerative lysosomal storage disorders involving ganglioside accumulation.^76^ Further investigation into the modifications and exact isomers of gangliosides in this abscess border would improve our understanding of ganglioside biology as it relates to foam cell development. Although macrophages seem to be the most well-studied in literature, we acknowledge our reported lipid distributions could still be associated with lipid-laden phenotypes of other cells like fibroblasts and neutrophils at the abscess border.

Finally, returning to the topic of inflammatory signaling, lipid droplets in the cytoplasm of foam cells are sources of arachidonic acid for eicosanoid production.^77^ Multiple studies have demonstrated pro-inflammatory or anti-inflammatory consequences of foam cell development that are largely dependent on the specific pathology and environmental conditions.^67,78–80^ Macrophages or foam cells in the context of a *S. aureus* abscess are not as well characterized compared to those in granulomas or atheromas, and any conclusion on the inflammatory function of the specialized cells in the fibrotic border of a *S. aureus* abscess cannot be supported with current data. Continued exploration of the cellular and molecular landscape in this region should reveal mechanisms of inflammation stimulation or resolution and foam cell development.

## Limitations of the Study

Although MALDI IMS supplemented with LC-MS/MS provides confirmation of lipid class/head group and fatty acid composition for lipids present in the whole organ, this workflow does not provide spatial distributions of isomeric species. Some lipids with multiple fatty acid combinations were identified with LC-MS/MS, but we did not employ a method to definitively match these isomers to its ion image. In this study, the ion images are composite maps of all isomeric species that were identified with LC-MS/MS, and we used the relative abundances between the two techniques to infer distributions. For example, [PC(38:6) + K]^+^ was the most abundant lipid in the muscle and was less abundant in the outer portion of the abscess from the imaging data. LC-MS/MS experiments yielded a large chromatographic peak for PC(16:0_22:6) and a substantially smaller peak area for PC(18:2_20:4). These identifications were paired to tissue types using relative abundances, but either isomer could be present in either tissue type. High throughput spatial MS/MS experiments and further ion mobility-based imaging are required to interrogate all isomer distributions.

Additionally, not all lipid classes could be validated with the bulk reversed-phase LC-MS/MS experiments. Bacterial lipids like PGs, Lysyl-PGs, and CLs were not abundant enough in homogenates for detection, but some have been confirmed by on-tissue MALDI MS/MS of *S. aureus* kidney abscesses.^28^ Chromatography and mass detection was not optimal for GMs; however, preliminary identifications of GM1, GM2, GM3, and respective derivatives of each (GalNAc- and +O modifications) are consistent with validated identifications of gangliosides in *S. aureus* kidney abscesses.^81^ Ether PAs, PAs, and CerPs were also not detected by LC-MS/MS even though a deuterated PA was included as an internal standard. These lipids were not retained suggesting further optimization of chromatography is necessary to confirm the identity of lipids with a free phosphate group.

Finally, our imaging platform and untargeted experimental design cannot definitively link lipid distributions to exact leukocyte populations, specifically at the abscess border. Multiplexed immunofluorescence combined with computational data mining tools must be employed to fully conclude lipids are direct markers of multiple cell-specific processes. Incorporating spatial proteomics and transcriptomics technologies in the future would also help confirm hypotheses and reveal new cellular factors driving these alterations in lipid metabolism.

## METHODS

### Chemicals

Acetone, ammonium formate (AF), carboxymethyl cellulose (CMC), glycerol, and 2,5-dihydroxyacetophenone (DHA, further purified by recrystallization) were purchased from Sigma-Aldrich (St. Louis, MO, USA). Isopentane, ethanol (EtOH), isopropanol (IPA), 10% neutral buffered formalin (NBF), and gelatin from Fisher Scientific (Pittsburgh, PA, USA). A Clearium mounting medium was purchased from Electron Microscopy Sciences (Hatfield, PA, USA). Ultrapure solvents and chemicals (LiChrosolv®) for lipid extractions and liquid chromatography, including methanol (MeOH), acetonitrile (ACN), IPA, H_2_O, methyl *tert*-butyl ether (MTBE), and AF were purchased from MilliporeSigma (Burlington, MA, USA). Ultrapure formic acid (FA) was purchased from Thermo Scientific (Rockford, IL, USA).

Harris hematoxylin (H) and eosin Y (E) were purchased from Thermo Scientific (Pittsburgh, PA, USA). For Masson’s Trichrome (MTC) staining, bouin’s fluid (BF), weigert’s hematoxylin (WH), biebrich scarlet-acid fuschsin (BSAF), phosphomolybdic/phosphotungstic acid (PPA), aniline blue stain (ABS), and acetic acid (AA) were purchased from American Mastertech Scientific Inc. (Lodi, CA, USA). For Oil Red O (ORO) staining, proprietary pre-stain and differentiation solution, oil red o (ORO), and mayer’s hematoxylin (MH) were purchased from VitroVivo Biotech (Rockville, MD, USA).

For immunohistochemistry reagents, peroxidase blocking solution (BLOXALL®), 3,3′-Diaminobenzidine (DAB; ImmPACT® DAB EqV), and modified MH were purchased from Vector Laboratories (Newark, CA, USA). Universal blocking reagent (Power Block™) was purchased from BioGenex (Femont, CA, USA). BOND primary antibody diluent was purchased from Leica Biosystems (Wetzlar, Germany). Primary rat anti-F4/80 IgG was purchased from Novusbio (location). Secondary rabbit anti-rat IgG and tertiary goat anti-rabbit IgG with horseradish peroxidase (HRP; ImmPRESS® HRP) was purchased from Vector Laboratories (Newark, CA, USA).

### Murine Model of *S. aureus* Osteomyelitis

*S. aureus* strain, USA300 AH1263 *attC*::P*sarA*_sfGFP, was used for all experiments (PMID: 3161596). AH1263 is a derivative of strain LAC that is erythromycin- and tetracycline-sensitive and represents a common *S. aureus* lineage that is isolated from musculoskeletal infections.^82^ Bacterial cultures were grown in 5 mL of tryptic soy broth (TSB) at 37°C with shaking at 180 rpm overnight. Methods for inducing osteomyelitis in mice were previously described in extensive detail.^39^ In summary, left femurs of seven- to nine-week-old female C57BL/6J mice (Jackson Laboratory, Bar Harbor, ME, USA) were exposed and a cortical bone defect was created in the diaphysis. A *S. aureus* inoculum (∼1 × 10^6^ CFU in 2 μL phosphate buffered saline (PBS)) was delivered via intraosseous injection through the femur defect. Mock-infected controls were constructed by creating a bone defect and delivering PBS instead of a bacterial inoculum. Mice were humanely euthanized 14 days post-infection, and infected and mock-infected femurs were excised with the majority of surrounding muscle being surgically removed. The femurs were immediately snap frozen over a dry ice-isopentane slurry and stored at -80 °C until further processing. For high spatial resolution imaging experiments, the only changes to the procedure were the use of Catchup mice (neutrophils expressing tdTomato), and the intact leg (femur, tibia, fibula, muscle) was excised.^44^

A cohort of 3 infected mice and 3 mock-infected mice were used for the main dataset involved in the majority of experimental analyses (**Figures 1-6**). Additional Catchup mice were used for the high spatial resolution imaging experiments (**Figure 7**). All animal handling and experimental procedures were approved by the Vanderbilt University Institutional Animal Care and Use Committee (IACUC) and performed in accordance with NIH guidelines, the Animal Welfare Act, and U.S. federal law.

### Sample Preparation

Methods for preparing fresh-frozen bone compatible with high spatial resolution MALDI IMS and microscopy were previously described in extensive detail.^38^ In summary, frozen femurs were embedded in gelatinous mixture of 5% CMC and 10% gelatin and refrozen. Femurs were cut at a cryosection thickness of 8 μm using a CM3050 S cryostat (Leica Biosystems, Wetzlar, Germany). Transparent Cryofilm 3C 16UF (SECTION-LAB, Hiroshima, Japan) was used in replace of an anti-roll bar and was necessary to retain the integrity and morphology of undecalcified bone tissue.^83^ A handheld fluorescence microscope, Dino-Lite AM4115T-GRFBY (AnMo Electronics, Taipei City, Taiwan), was used to detect the bacterial fluorophore while cryosectioning to ensure every collected section contained a SAC **(Figure S1).** The section bound to Cryofilm was mounted to conductive indium tin oxide (ITO)-coated glass slides (Delta Technologies, Loveland, CO, USA) using transparent ZIG 2 Way Glue (Kuretake Co, Nara, Japan). One infected and one mock-infected section were mounted onto the same slide and were subsequently stored at -80 °C. On the day of the MALDI IMS experiments, frozen samples were freeze-dried for 1.5 h using a glass apparatus kept at ambient temperature and ∼150-300 mTorr. For histological purposes, adjacent (not truly serial) sections were also collected, fixed, and stained following the protocols below rather than freeze-dried. For the salt washing experiment, a freeze-dried sample was washed with two cold solutions of 150 mM AF for 20 s each, and subsequently dried with nitrogen gas.

Fluorescence images of femur sections were collected prior to matrix application. 2 mL of a 20 mg/mL solution of DHA in acetone was dispersed onto a heated surface and sublimed onto sample slides using an in-house sublimation apparatus. This resulted in a uniform matrix layer with a surface density of 2.6 μg/mm^2^. The deposited matrix was recrystallized using 1 mL of 5% IPA in a hydration chamber set at 55 °C for 1.5 min. Following MALDI IMS, sections were washed, stained, and imaged for histological annotations.

For the main dataset, three sections (section replicates) for each of three infected femurs and three mock-infected femurs (biological-femur replicates) were prepared, yielding 18 femur sections on nine sample slides. Groups of three samples, each from a different biological cohort were prepared for imaging together to delineate sample preparation and instrumental variation from biological variation; in other words, all section replicates for a single femur were not prepared and run together.

### MALDI IMS Acquisition

All MALDI IMS data were acquired using a timsTOF fleX (Bruker Daltonics, Bremen, Germany).^84^ All imaging experiments were operated in qTOF mode with TIMS deactivated unless otherwise noted. For acquisition of the main dataset, a pitch offset strategy was employed. The SmartBeam 3D 10 kHz frequency tripled Nd:YAG laser (355 nm) was focused and beam scan activated to produce a 10 μm x 10 μm burn pattern. The z position of the stage was adjusted to keep the laser focused and account for additional height of the mount and Cryofilm. Positive ion data were acquired first with a stage pitch of 20 μm. Stage coordinates in Fleximaging were then offset 10 μm in each x-y dimension, and negative ion data were acquired with the same 20 μm stage pitch. This strategy yields an effective spatial resolution of 20 μm for both polarities from the same pixel regions. The laser was set to 80% power (0% attenuator offset) and 100 shots. For high spatial resolution (10 μm) experiments, beam scan was still implemented, but the stage pitch was set to 10 μm. Prior to data acquisition in positive and negative ion modes, ESI-L Tune Mix (Agilent Technologies, Santa Clara, CA, USA) was infused into the system for external mass calibration. For both polarities, data were acquired from *m/*z 150-2000. For positive ion mode, the following MS1 parameters were set: <u>Transfer</u>-MALDI Plate Offset= 30.0 V, Deflection 1 Delta= 70.0 V, Funnel 1 RF= 450.0 Vpp, isCID Energy= 0.0 eV, Funnel 2 RF= 500.0 Vpp, Multipole RF= 500.0 Vpp; <u>Collision Cell</u>- Collision Energy= 10.0 eV, Collison RF= 2900.0 Vpp; <u>Quadrupole</u>- Ion Energy= 5.0 eV, Low Mass= *m/z* 300.00; <u>Focus Pre TOF</u>- Transfer Time= 110.0 µs, Pre Pulse Storage= 10.0 µs. For negative ion mode, changes include: MALDI Plate Offset= -30.0 V, Deflection 1 Delta= -70.0 V, Collision Energy= -10.0 eV, Ion Energy= -5.0 eV.

For the TIMS imaging experiment (**Figure S7**), negative ion data were acquired on a prototype timsTOF fleX with TIMS activated. Beam scan was used to produce a 20 μm x 20 μm burn pattern and the stage pitch was set to 20 μm. The laser was adjusted to 70% power (0% attenuator offset) and 200 shots. The following MS1 parameters were set: MS range= *m/z* 400-2000; <u>Transfer</u>- MALDI Plate Offset= -70.0 V, Deflection 1 Delta= -70.0 V, Funnel 1 RF= 350.0 Vpp, isCID Energy= -10.0 eV, Funnel 2 RF= 400.0 Vpp, Multipole RF= 500.0 Vpp; <u>Collision Cell</u>- Collision Energy= -15.0 eV, Collison RF= 2500.0 Vpp; <u>Quadrupole</u>- Ion Energy= -10.0 eV, Low Mass= *m/z* 500.00; <u>Focus Pre TOF</u>- Transfer Time= 70.0 µs, Pre Pulse Storage= 12.0 µs. The following TIMS parameters were set: 1/K_0_ range= 1.10-1.70 Vs/cm^2^, Ramp Time= 400.0 ms, Accumulation Time= 20.1 ms; <u>Offsets</u>- Δt1= 20.0 V, Δt2= 120.0 V, Δt3= -70.0 V, Δt4= -100.0 V, Δt5= 0.0 V, Δt6= -100.0 V, Collision Cell In= -300.0 V.

### Histology

H&E, MTC, and ORO staining were performed on a different section replicate from each femur following MALDI IMS **(Figure S1).** For all post-IMS stains, DHA was removed using an ethanol wash prior to beginning the H&E and MTC protocols and sublimed off prior to the ORO protocol. Protocols are established from common practices and kits, but key highlights are as followed; for H&E, sections were stained sequentially with H for 2 min 15 s and E for 45 s; for MTC, sections were fixed with bouins fluid at ambient temperature overnight and stained sequentially with WH for 5 min, BSAF for 15 min, PPA for 15 min, ABS for 10 min, and 1% AA for 3 min; for ORO, sections were fixed with NBF for 20 min and stained sequentially with ORO for 10 minutes and MH for 1 min. Coverslips were mounted on stained femurs using Clearium mounting medium for H&E and MTC and 50:50 glycerol:H_2_O for ORO. Adjacent sections collected for cellular analysis with no bone marrow cracks were thawed for 10 s from a frozen state and fixed using 10 s of ethanol and 2 min of NBF.^85^ Most importantly, the sections remained in liquid throughout the staining protocols until coverslips were in place. Histopathology was performed by a Board-Certified Veterinary Pathologist.

For F4/80 IHC, sections were thawed for 10 s from a frozen state and fixed using 10 s of ethanol and 5 min of NBF. With PBS or H_2_O washes between most steps, sections were sequentially exposed to peroxidase blocking solution for 30 min, universal blocking reagent for 30 min, 1:600 primary anti-F4/80 for 1 h, 1:2k secondary anti-rat for 15 min, tertiary anti-rabbit HRP for 10 min, DAB for 10 min, and modified MH for 45 s. Femur sections following the same protocol without primary antibody incubation served as a control and possessed no DAB staining.

### Microscopy

An AxioScan.Z1 slide scanner (Carl Zeiss Microscopy GmbH, Oberkochen, Germany) was utilized for fluorescence and bright-field microscopy. Fluorescence images were collected using a 10x microscope objective and EGFP (ex: 488 nm, em: 509 nm), DAPI (ex: 353 nm, em: 465 nm), and DsRed (ex: 545 nm, em: 572 nm) filters. Unfiltered LEDs were used to acquire 20x bright-field images. To compensate for height differences across the tissue surface, a z-stack of five depths over a 35 μm range was acquired at every tile and merged using extended depth of focus in the Zen software suite (Carl Zeiss Microscopy GmbH, Oberkochen, Germany). For fluorescence images, false colors were assigned to each channel (EGFP= green, DAPI= blue, DsRed= orange). Unless otherwise noted, intensity scaling for each fluorescence channel was optimized and applied consistently to all images of the main dataset **(Figure S1).** All bright-field images were white balanced and consistently saturated in Illustrator CS6 (Adobe Inc., San Jose, CA, USA) to enhance stain contrast unless otherwise noted.

### Lipid Extraction for LC-MS/MS

Methods for extracting lipids from small amounts of tissue were described in detail in a protocols.io.^86^ In summary, 40 µm of cryosection tissue that was adjacent to the imaging sections for each femur was collected into a glass vial. Methanol, Equisplash® internal standard (Avanti Polar Lipids, Alabaster, AL, USA), and metal beads were added. Sections were homogenized through a combination of vortexing, dry/wet ice exposure, and sonication. Sample vials were dried under N_2_ and lipids were subsequently extracted using 4:1:1 MTBE:MeOH:H_2_O, which was modified from an existing protocol.^87^ Following proper mixing and phase separation, a majority of the MTBE layer was collected, dried, and reconstituted in MeOH for LC-MS/MS analysis.

### PASEF LC-MS/MS

An ACQUITY Premier UPLC (Waters Corporation, Milford, MA, USA) with a Premier C18 CSH column (2.1 mm x 100 mm, Waters Corporation) was connected to the Apollo electrospray source of the timsTOF fleX. Sample volumes of 5 µL for positive ion mode and 10 µL for negative ion mode were injected and reversed phase separation was performed over a 35 min gradient using 60:40 ACN:H_2_O-10 mM AF-0.1% FA (mobile phase A) and 90:10 IPA:ACN-10 mM AF-0.1% FA (mobile phase B) at a 200 mL/min flow rate and 60 °C column temperature.

Parallel accumulation-serial fragmentation (PASEF) was employed to obtain quality MS/MS spectra for lipids.^88^ With TIMS on, the system was calibrated to 132.5 V for the positive ion, *m/*z 622.029 in the ESI-L Tune Mix, and 118.5 V for the negative ion, *m/z* 601.979, by altering the tunnel-in pressure. For positive ion mode (and negative ion mode), the following MS1 parameters were set: MS range= *m/z* 50-2000; <u>Transfer</u>- Deflection 1 Delta= 80.0 (-80.0) V, Funnel 1 RF= 500.0 (250.0) Vpp, isCID Energy= 0.0 eV, Funnel 2 RF= 300.0 Vpp, Multipole RF= 250.0 Vpp; <u>Collision Cell</u>- Collision Energy= 10.0 (-10.0) eV, Collison RF= 1800.0 Vpp; <u>Quadrupole</u>- Ion Energy= 5.0 (-5.0) eV, Low Mass= m/z 50.00; <u>Focus Pre TOF</u>- Transfer Time=

65.0 (70.0) µs, Pre Pulse Storage= 5.0 (10.0) µs. The following TIMS parameters were set: 1/K_0_ range= 0.45-1.90 (0.45-1.88) Vs/cm^2^, Ramp Time= 100.0 ms, Accumulation Time= 100.0 ms; <u>Offsets</u>- Δt1= -20.0 (20.0) V, Δt2= -120.0 (120.0) V, Δt3= 80.0 (-80.0) V, Δt4= 100.0 (-100.0) V, Δt5= 0.0 V, Δt6= 150.0 (-100.0) V, Collision Cell In= 250.0 (-250.0) V. The following MS/MS parameters were set: <u>Scan 1</u>- Collision Energy= 30.0 (-40.0) eV at 1/K_0_ of 0.45 (0.55) and 30.0 (-40.0) eV at 1/K_0_ of 1.90 (1.88), Collision RF= 450.0 Vpp, Transfer Time= 25.0 µs, Pre Pulse Storage= 5.0 µs; <u>Scan 2</u>- Collision Energy= 30.0 (-40.0) eV at 1/K_0_ of 0.45 (0.55) and 50.0 (-65.0) eV at 1/K_0_ of 1.90 (1.88), Collision RF= 1800.0 Vpp, Transfer Time= 75.0 (90.0) µs, Pre Pulse Storage= 10.0 µs. Data analysis is performed in MS-DIAL v4.90 which automatically searches a lipid database and assigns annotations based on MS/MS dot product, reverse dot product, mobility information, and retention time.^89^ For these annotations, every chromatographic peak and corresponding MS/MS spectra were manually inspected and confirmed.

### Lipid Identifications

Lipids were preliminarily annotated in MALDI IMS data using accurate mass with a mass error tolerance of 3 ppm compared to the LIPIDMAPS database (lipidmaps.org).^90,91^ For some lower abundant lipids, data were manually searched with the aid of LipidPioneer.^92^ To help confirm preliminary identifications, ionizing adducts (e.g. H^+^, Na^+^, K^+^, and H^-^) were corroborated with experiments using AF washes; salt adducts decrease and pronated species increase following the wash. Preliminary identifications were supplemented with LC-MS/MS confirmed identifications where possible. Relative retention time and inverse mobility also helped increase confidence. Primary identifications for a given *m/z* were assigned to the most abundant peak of the extracted ion chromatogram (0.01 Da MS1 tolerance). Secondary identifications were assigned to smaller chromatographic peaks and some less intense fragments in MS/MS spectra that were not separated by chromatography. Some lipids could not be confirmed by MS/MS, most likely due to their low abundance within the tissue. Whole lipid classes that have no MS/MS confirmatory evidence (i.e. PAs, CerPs) are marked as lower confidence. In general, large charts, plots, and maps are labeled with preliminary identifications, and ion images are labeled with primary identifications, unless otherwise noted.

### Imaging Based Analyses

MALDI IMS data were visualized, and imaging analyses were conducted using SCiLS Lab v2023 (Bruker Daltonics). Root mean square normalization was applied for all analyses including ion image visualization. The selection window for all ion images was set from 15-20 ppm depending on its proximity to a nearby peak. Image intensity scales were manually set for optimal pixel contrast, unless otherwise noted. For untargeted investigation of the murine lipidome, average mass spectra from each femur were peak picked using the sliding window function and consolidated into one list. These *m/z* features were analyzed by *k*-means clustering where clustering was completed on the combination of all section replicates from each infected and mock-infected femur using Euclidean distance and no denoising. Analysis was also performed for each polarity, separately. The number of clusters was varied, and the optimal number was determined qualitatively by assessing the resulting segmented images. Ultimately, *k*=13 was used for positive ion mode and *k*=11 for negative ion mode. Note, positive and negative ion mode clustering results are referred to as +*k* and -*k*, respectively. Both cases provided the most cluster segments that deviated from background noise (signal from matrix, tape, etc.), while still being biologically relevant. A lower *k* led to a loss of segments that marked known tissue types, and a higher *k* started to segment out small deviations in known tissue types that had questionable biological relevance. For example, the positive cluster marking the muscle split into two (+*k*5 and +*k*6) and the negative cluster marking the physis split into two (-*k*5 and -*k*6); the physis segments were given the same color because the pixel area was too small to be visualized. This initial splitting was kept in the analysis to show these small deviations begin to form. Top lipid markers for a cluster or tissue type were identified by dividing the lipid’s mean intensity within a cluster over its mean intensity across the whole section, yielding a cluster ratio. Mean intensity values were extracted from the maximum peak value in the average mass spectrum of a given region. All identified lipids above an arbitrary ratio of 2 for each cluster are reported. If a known tissue type, like the nerve, was not properly segmented by *k*-means clustering, the corresponding pixels were manually extracted using contrasting ion images and analyzed in the same manner.

For targeted investigation of the *S. aureus* lipidome, a fluorescence image collected prior to MALDI IMS was registered to IMS data using the SCiLS registration function, and regions of interest were manually drawn around the fluorescent SACs. For targeted investigation of the abscess, *k*-means clustering was performed on all abscesses simultaneously using the selected four ions, Euclidian distance, weak denoising, and a *k* of 5 for each polarity. Using identifications from the previous tissue marker analysis or manually screening for new lipids, each lipid’s mean intensity for the outside abscess cluster was compared to its mean intensity for the inside abscess cluster. A paired parametric t-test was performed to test significant changes between the clusters. For analysis of mock-infected tissue, the clusters associated with the bone marrow for positive ion mode (+*k*3, +*k*4) were combined, termed BMComb, and clusters for negative ion mode (-*k*1, -*k*2, -*k*3) were similarly combined. For each lipid identified in the bone marrow, its mean intensity for infected BMComb was compared to its mean intensity for mock-infected BMComb. Although every pixel might not associate with the bone marrow, this approach was more representative than comparing the full sections that include unrelated tissue types and off-tissue background noise. Lipid distributions associated with the abscess analysis were then analyzed using PCA, which was performed on all nine infected BMComb simultaneously for each polarity. Each pixel within the entire cohort of samples was treated as an observation. Observations were then reduced to an interpretable loadings plots, where every loading is the distribution of a single lipid across the cohort. For the abscess border analysis, a Pearson’s correlation was performed on a region encompassing a single large abscess in the intramedullary cavity. Coefficients were calculated between an ion’s distribution in this region and the ring-like distributions of [PC O-(16:0_20:4) + K]^+^ for positive ion mode and [PI O-(18:1_20:4) - H]^-^ for negative ion mode. New lipids were identified as a result of this analysis, and these lipids along with the previous abscess lipids were incorporated into the heat maps. All bar graphs and heat maps were created using Prism (GraphPad.Software, San Diego, CA, USA).

## SIGNIFICANCE

MALDI IMS and supporting microscopy were leveraged to provide *in situ* lipid distributions for various tissue types throughout *S. aureus*-infected femurs. These multiplexed and spatially resolved lipidomics data offer a unique perspective into cellular dynamics that *in vitro* experiments would struggle to recapitulate due to the complex cellular factors and tissue microenvironments driving abscess pathology. Molecular imaging also informs the phenotype of a subpopulation of cells, in this case, macrophages, that cannot be discerned with traditional microscopy modalities. Specific species belonging to classes of ether lipids and arachidonoyl lipids were closely associated with leukocytes and the formation of bone marrow abscesses. The lipids reported at the abscess border provided chemical insight into what could be various cellular processes of fibrosis, inflammatory signaling, lipid metabolism dysregulation, and lysosomal dysfunction. Given that this model represents a late-stage infection, cellular and molecular information is valuable to understanding why this infection progresses and evades immune clearance. Finally, while the biological focus of this work is *S. aureus* osteomyelitis, this imaging platform can be applied to other models to understand complex inflammatory and metabolic disorders that compromise bone biology and hematopoiesis.

## AUTHOR CONTRIBUTIONS

The manuscript was written through contributions of all authors. All authors have given approval to the final version of the manuscript.

## NOTES

The authors declare no competing financial interest.

## ACKNOWLEDGEMENTS

The authors would like to thank Dr. Eric Spivey and Dr. David Anderson for creating the in-house sublimation apparatus, Cindy Lowe for her IHC guidance, and Dr. Elizabeth Neumann for her professional mentorship while at VU. Support was provided by the NIH National Institute of Allergy and Infectious Disease (R01 AI145992 awarded to J.M.S. and J.E.C., R01 AI161022 and R01 AI173795 awarded to J.E.C, and R01AI138581 awarded to J.M.S.), the NIH Shared Instrumentation Grant Program (S10 OD012359 awarded to R.M.C.), and the National Science Foundation Major Research Instrument Program (CBET 1828299 awarded to J.M.S. and R.M.C.). We acknowledge the Translational Pathology Shared Resource supported by the NCI/NIH Cancer Center Support Grant P30CA068485. C.J.G. and C.E.B. were supported by the NIH National Institute of Allergy and Infectious Disease (T32 AI112541). J.E.C. is supported by the NIH National Institute of Allergy and Infectious Disease (R01 AI132560) and a Burroughs Welcome Fund Career Award for Medical Scientists.

## Notes

### Competing Interest Statement

The authors have declared no competing interest.

## REFERENCES

1. Gimza, B. D.; Cassat, J. E. Mechanisms of Antibiotic Failure During Staphylococcus Aureus Osteomyelitis. Front. Immunol. 2021, 12, 303. 10.3389/fimmu.2021.638085.

2. Conterno, L. O.; Turchi, M. D. Antibiotics for Treating Chronic Osteomyelitis in Adults. Cochrane Database Syst. Rev. 2013, No. 9, CD004439. 10.1002/14651858.CD004439.pub3.

3. Kremers, H. M.; Nwojo, M. E.; Ransom, J. E.; Wood-Wentz, C. M.; Melton, L. J.; Huddleston, P. M. Trends in the Epidemiology of Osteomyelitis. J. Bone Joint Surg. Am. 2015, 97 (10), 837–845. 10.2106/JBJS.N.01350.

4. Lew, D. P.; Waldvogel, F. A. Osteomyelitis. The Lancet 2004, 364 (9431), 369–379. 10.1016/S0140-6736(04)16727-5.

5. Kavanagh, N.; Ryan, E. J.; Widaa, A.; Sexton, G.; Fennell, J.; O’Rourke, S.; Cahill, K. C.; Kearney, C. J.; O’Brien, F. J.; Kerrigan, S. W. Staphylococcal Osteomyelitis: Disease Progression, Treatment Challenges, and Future Directions. Clin. Microbiol. Rev. 2018, 31 (2). 10.1128/CMR.00084-17.

6. Walter, G.; Kemmerer, M.; Kappler, C.; Hoffmann, R. Treatment Algorithms for Chronic Osteomyelitis. Dtsch. Ärztebl. Int. 2012, 109 (14), 257–264. 10.3238/arztebl.2012.0257.

7. Lowy, F. D. Staphylococcus Aureus Infections. N. Engl. J. Med. 1998, 339 (8), 520–532. 10.1056/NEJM199808203390806.

8. Cheng, A. G.; DeDent, A. C.; Schneewind, O.; Missiakas, D. A Play in Four Acts: Staphylococcus Aureus Abscess Formation. Trends Microbiol. 2011, 19 (5), 225–232. 10.1016/j.tim.2011.01.007.

9. Peschel, A.; Jack, R. W.; Otto, M.; Collins, L. V.; Staubitz, P.; Nicholson, G.; Kalbacher, H.; Nieuwenhuizen, W. F.; Jung, G.; Tarkowski, A.; van Kessel, K. P.; van Strijp, J. A. Staphylococcus Aureus Resistance to Human Defensins and Evasion of Neutrophil Killing via the Novel Virulence Factor MprF Is Based on Modification of Membrane Lipids with L-Lysine. J. Exp. Med. 2001, 193 (9), 1067–1076. 10.1084/jem.193.9.1067.

10. Ernst, C. M.; Peschel, A. Broad-Spectrum Antimicrobial Peptide Resistance by MprF-Mediated Aminoacylation and Flipping of Phospholipids. Mol. Microbiol. 2011, 80 (2), 290–299. 10.1111/j.1365-2958.2011.07576.x.

11. Braverman, N. E.; Moser, A. B. Functions of Plasmalogen Lipids in Health and Disease. Biochim. Biophys. Acta BBA - Mol. Basis Dis. 2012, 1822 (9), 1442–1452. 10.1016/j.bbadis.2012.05.008.

12. Dean, J. M.; Lodhi, I. J. Structural and Functional Roles of Ether Lipids. Protein Cell 2018, 9 (2), 196–206. 10.1007/s13238-017-0423-5.

13. Rangholia, N.; Leisner, T. M.; Holly, S. P. Bioactive Ether Lipids: Primordial Modulators of Cellular Signaling. Metabolites 2021, 11 (1), 41. 10.3390/metabo11010041.

14. Jiménez-Rojo, N.; Riezman, H. On the Road to Unraveling the Molecular Functions of Ether Lipids. FEBS Lett. 2019, 593 (17), 2378–2389. 10.1002/1873-3468.13465.

15. Roy Baker, R. The Eicosanoids: A Historical Overview. Clin. Biochem. 1990, 23 (5), 455–458. 10.1016/0009-9120(90)90255-S.

16. Wang, B.; Wu, L.; Chen, J.; Dong, L.; Chen, C.; Wen, Z.; Hu, J.; Fleming, I.; Wang, D. W. Metabolism Pathways of Arachidonic Acids: Mechanisms and Potential Therapeutic Targets. Signal Transduct. Target. Ther. 2021, 6 (1), 1–30. 10.1038/s41392-020-00443-w.

17. Caprioli, R. M.; Farmer, T. B.; Gile, J. Molecular Imaging of Biological Samples: Localization of Peptides and Proteins Using MALDI-TOF MS. Anal. Chem. 1997, 69 (23), 4751–4760. 10.1021/ac970888i.

18. Cornett, D. S.; Reyzer, M. L.; Chaurand, P.; Caprioli, R. M. MALDI Imaging Mass Spectrometry: Molecular Snapshots of Biochemical Systems. Nat. Methods 2007, 4 (10), 828–833. 10.1038/nmeth1094.

19. Chaurand, P.; Schwartz, S. A.; Billheimer, D.; Xu, B. J.; Crecelius, A.; Caprioli, R. M. Integrating Histology and Imaging Mass Spectrometry. Anal. Chem. 2004, 76 (4), 1145–1155. 10.1021/ac0351264.

20. Abdelmoula, W. M.; Škrášková, K.; Balluff, B.; Carreira, R. J.; Tolner, E. A.; Lelieveldt, B. P. F.; van der Maaten, L.; Morreau, H.; van den Maagdenberg, A. M. J. M.; Heeren, R. M. A.; McDonnell, L. A.; Dijkstra, J. Automatic Generic Registration of Mass Spectrometry Imaging Data to Histology Using Nonlinear Stochastic Embedding. Anal. Chem. 2014, 86 (18), 9204–9211. 10.1021/ac502170f.

21. Patterson, N. H.; Tuck, M.; Van de Plas, R.; Caprioli, R. M. Advanced Registration and Analysis of MALDI Imaging Mass Spectrometry Measurements through Autofluorescence Microscopy. Anal. Chem. 2018, 90 (21), 12395–12403. 10.1021/acs.analchem.8b02884.

22. Van de Plas, R.; Yang, J.; Spraggins, J.; Caprioli, R. M. Image Fusion of Mass Spectrometry and Microscopy: A Multimodality Paradigm for Molecular Tissue Mapping. Nat. Methods 2015, 12 (4), 366–372. 10.1038/nmeth.3296.

23. Corbin, B. D.; Seeley, E. H.; Raab, A.; Feldmann, J.; Miller, M. R.; Torres, V. J.; Anderson, K. L.; Dattilo, B. M.; Dunman, P. M.; Gerads, R.; Caprioli, R. M.; Nacken, W.; Chazin, W. J.; Skaar, E. P. Metal Chelation and Inhibition of Bacterial Growth in Tissue Abscesses. Science 2008, 319 (5865), 962–965. 10.1126/science.1152449.

24. Attia, A. S.; Schroeder, K. A.; Seeley, E. H.; Wilson, K. J.; Hammer, N. D.; Colvin, D. C.; Manier, M. L.; Nicklay, J. J.; Rose, K. L.; Gore, J. C.; Caprioli, R. M.; Skaar, E. P. Monitoring the Inflammatory Response to Infection through the Integration of MALDI IMS and MRI. Cell Host Microbe 2012, 11 (6), 664–673. 10.1016/j.chom.2012.04.018.

25. Spraggins, J. M.; Rizzo, D. G.; Moore, J. L.; Rose, K. L.; Hammer, N. D.; Skaar, E. P.; Caprioli, R. M. MALDI FTICR IMS of Intact Proteins: Using Mass Accuracy to Link Protein Images with Proteomics Data. J. Am. Soc. Mass Spectrom. 2015, 26 (6), 974–985. 10.1021/jasms.8b05050.

26. Cassat, J. E.; Moore, J. L.; Wilson, K. J.; Stark, Z.; Prentice, B. M.; Van de Plas, R.; Perry, W. J.; Zhang, Y.; Virostko, J.; Colvin, D. C.; Rose, K. L.; Judd, A. M.; Reyzer, M. L.; Spraggins, J. M.; Grunenwald, C. M.; Gore, J. C.; Caprioli, R. M.; Skaar, E. P. Integrated Molecular Imaging Reveals Tissue Heterogeneity Driving Host-Pathogen Interactions. Sci. Transl. Med. 2018, 10 (432), eaan6361. 10.1126/scitranslmed.aan6361.

27. Perry, W. J.; Spraggins, J. M.; Sheldon, J. R.; Grunenwald, C. M.; Heinrichs, D. E.; Cassat, J. E.; Skaar, E. P.; Caprioli, R. M. Staphylococcus Aureus Exhibits Heterogeneous Siderophore Production within the Vertebrate Host. Proc. Natl. Acad. Sci. 2019, 116 (44), 21980–21982. 10.1073/pnas.1913991116.

28. Perry, W. J.; Grunenwald, C. M.; Van De Plas, R.; Witten, J. C.; Martin, D. R.; Apte, S. S.; Cassat, J. E.; Pettersson, G. B.; Caprioli, R. M.; Skaar, E. P.; Spraggins, J. M. Visualizing Staphylococcus Aureus Pathogenic Membrane Modification within the Host Infection Environment by Multimodal Imaging Mass Spectrometry. Cell Chem. Biol. 2022, 29 (7), 1209–1217.e4. 10.1016/j.chembiol.2022.05.004.

29. Sharman, K.; Patterson, N. H.; Migas, L. G.; Neumann, E. K.; Allen, J.; Gibson-Corley, K. N.; Spraggins, J. M.; Van de Plas, R.; Skaar, E. P.; Caprioli, R. M. MALDI IMS-Derived Molecular Contour Maps: Augmenting Histology Whole-Slide Images. J. Am. Soc. Mass Spectrom. 2023, 34 (5), 905–912. 10.1021/jasms.2c00370.

30. Hood, M. I.; Mortensen, B. L.; Moore, J. L.; Zhang, Y.; Kehl-Fie, T. E.; Sugitani, N.; Chazin, W. J.; Caprioli, R. M.; Skaar, E. P. Identification of an Acinetobacter Baumannii Zinc Acquisition System That Facilitates Resistance to Calprotectin-Mediated Zinc Sequestration. PLOS Pathog. 2012, 8 (12), e1003068. 10.1371/journal.ppat.1003068.

31. Moore, J. L.; Becker, K. W.; Nicklay, J. J.; Boyd, K. L.; Skaar, E. P.; Caprioli, R. M. Imaging Mass Spectrometry for Assessing Temporal Proteomics: Analysis of Calprotectin in Acinetobacter Baumannii Pulmonary Infection. PROTEOMICS 2014, 14 (7– 8), 820–828. 10.1002/pmic.201300046.

32. Zackular, J. P.; Moore, J. L.; Jordan, A. T.; Juttukonda, L. J.; Noto, M. J.; Nicholson, M. R.; Crews, J. D.; Semler, M. W.; Zhang, Y.; Ware, L. B.; Washington, M. K.; Chazin, W. J.; Caprioli, R. M.; Skaar, E. P. Dietary Zinc Alters the Microbiota and Decreases Resistance to Clostridium Difficile Infection. Nat. Med. 2016, 22 (11), 1330–1334. 10.1038/nm.4174.

33. Wexler, A. G.; Guiberson, E. R.; Beavers, W. N.; Shupe, J. A.; Washington, M. K.; Lacy, D. B.; Caprioli, R. M.; Spraggins, J. M.; Skaar, E. P. Clostridioides Difficile Infection Induces a Rapid Influx of Bile Acids into the Gut during Colonization of the Host. Cell Rep. 2021, 36 (10), 109683. 10.1016/j.celrep.2021.109683.

34. Prideaux, B.; ElNaggar, M. S.; Zimmerman, M.; Wiseman, J. M.; Li, X.; Dartois, V. Mass Spectrometry Imaging of Levofloxacin Distribution in TB-Infected Pulmonary Lesions by MALDI-MSI and Continuous Liquid Microjunction Surface Sampling. Int. J. Mass Spectrom. 2015, 377, 699–708. 10.1016/j.ijms.2014.08.024.

35. Blanc, L.; Lenaerts, A.; Dartois, V.; Prideaux, B. Visualization of Mycobacterial Biomarkers and Tuberculosis Drugs in Infected Tissue by MALDI-MS Imaging. Anal. Chem. 2018, 90 (10), 6275–6282. 10.1021/acs.analchem.8b00985.

36. Hulme, H. E.; Meikle, L. M.; Wessel, H.; Strittmatter, N.; Swales, J.; Thomson, C.; Nilsson, A.; Nibbs, R. J. B.; Milling, S.; Andren, P. E.; Mackay, C. L.; Dexter, A.; Bunch, J.; Goodwin, R. J. A.; Burchmore, R.; Wall, D. M. Mass Spectrometry Imaging Identifies Palmitoylcarnitine as an Immunological Mediator during Salmonella Typhimurium Infection. Sci. Rep. 2017, 7 (1), 1– 10.1038/s41598-017-03100-5.

37. Yang, H.; Chandler, C. E.; Jackson, S. N.; Woods, A. S.; Goodlett, D. R.; Ernst, R. K.; Scott, A. J. On-Tissue Derivatization of Lipopolysaccharide for Detection of Lipid A Using MALDI-MSI. Anal. Chem. 2020, 92 (20), 13667–13671. 10.1021/acs.analchem.0c02566.

38. Good, C. J.; Neumann, E. K.; Butrico, C. E.; Cassat, J. E.; Caprioli, R. M.; Spraggins, J. M. High Spatial Resolution MALDI Imaging Mass Spectrometry of Fresh-Frozen Bone. Anal. Chem. 2022, 94 (7), 3165–3172. 10.1021/acs.analchem.1c04604.

39. Cassat, J. E.; Hammer, N. D.; Campbell, J. P.; Benson, M. A.; Perrien, D. S.; Mrak, L. N.; Smeltzer, M. S.; Torres, V. J.; Skaar, E. P. A Secreted Bacterial Protease Tailors the Staphylococcus Aureus Virulence Repertoire to Modulate Bone Remodeling during Osteomyelitis. Cell Host Microbe 2013, 13 (6), 759–772. 10.1016/j.chom.2013.05.003.

40. Bahney, C. S.; Zondervan, R. L.; Allison, P.; Theologis, A.; Ashley, J. W.; Ahn, J.; Miclau, T.; Marcucio, R. S.; Hankenson, K. D. Cellular Biology of Fracture Healing. J. Orthop. Res. Off. Publ. Orthop. Res. Soc. 2019, 37 (1), 35–50. 10.1002/jor.24170.

41. White, D. C.; Frerman, F. E. Extraction, Characterization, and Cellular Localization of the Lipids of Staphylococcus Aureus. J. Bacteriol. 1967, 94 (6), 1854–1867. 10.1128/jb.94.6.1854-1867.1967.

42. Hewelt-Belka, W.; Nakonieczna, J.; Belka, M.; Bączek, T.; Namieśnik, J.; Kot-Wasik, A. Untargeted Lipidomics Reveals Differences in the Lipid Pattern among Clinical Isolates of Staphylococcus Aureus Resistant and Sensitive to Antibiotics. J. Proteome Res. 2016, 15 (3), 914–922. 10.1021/acs.jproteome.5b00915.

43. Hines, K. M.; Alvarado, G.; Chen, X.; Gatto, C.; Pokorny, A.; Alonzo, F.; Wilkinson, B. J.; Xu, L. Lipidomic and Ultrastructural Characterization of the Cell Envelope of Staphylococcus Aureus Grown in the Presence of Human Serum. mSphere 2020, 5 (3), e00339–20. 10.1128/mSphere.00339-20.

44. Hasenberg, A.; Hasenberg, M.; Männ, L.; Neumann, F.; Borkenstein, L.; Stecher, M.; Kraus, A.; Engel, D. R.; Klingberg, A.; Seddigh, P.; Abdullah, Z.; Klebow, S.; Engelmann, S.; Reinhold, A.; Brandau, S.; Seeling, M.; Waisman, A.; Schraven, B.; Göthert, J. R.; Nimmerjahn, F.; Gunzer, M. Catchup: A Mouse Model for Imaging-Based Tracking and Modulation of Neutrophil Granulocytes. Nat. Methods 2015, 12 (5), 445–452. 10.1038/nmeth.3322.

45. Putnam, N. E.; Fulbright, L. E.; Curry, J. M.; Ford, C. A.; Petronglo, J. R.; Hendrix, A. S.; Cassat, J. E. MyD88 and IL-1R Signaling Drive Antibacterial Immunity and Osteoclast-Driven Bone Loss during Staphylococcus Aureus Osteomyelitis. PLoS Pathog. 2019, 15 (4). 10.1371/journal.ppat.1007744.

46. Vandenbosch, M.; Nauta, S. P.; Svirkova, A.; Poeze, M.; Heeren, R. M. A.; Siegel, T. P.; Cuypers, E.; Marchetti-Deschmann, M. Sample Preparation of Bone Tissue for MALDI-MSI for Forensic and (Pre)Clinical Applications. Anal. Bioanal. Chem. 2020. 10.1007/s00216-020-02920-1.

47. Cheng, S.-H.; Groseclose, M. R.; Mininger, C.; Bergstrom, M.; Zhang, L.; Lenhard, S. C.; Skedzielewski, T.; Kelley, Z. D.; Comroe, D.; Hong, H.; Cui, H.; Hoover, J. L.; Rittenhouse, S.; Castellino, S.; Jucker, B. M.; Alsaid, H. Multimodal Imaging Distribution Assessment of a Liposomal Antibiotic in an Infectious Disease Model. J. Control. Release Off. J. Control. Release Soc. 2022, 352, 199–210. 10.1016/j.jconrel.2022.08.061.

48. Mueller, H. W.; O’Flaherty, J. T.; Greene, D. G.; Samuel, M. P.; Wykle, R. L. 1-O-Alkyl-Linked Glycerophospholipids of Human Neutrophils: Distribution of Arachidonate and Other Acyl Residues in the Ether-Linked and Diacyl Species. J. Lipid Res. 1984, 25 (4), 383–388. 10.1016/S0022-2275(20)37812-3.

49. Ivanova, P. T.; Milne, S. B.; Brown, H. A. Identification of Atypical Ether-Linked Glycerophospholipid Species in Macrophages by Mass Spectrometry. J. Lipid Res. 2010, 51 (6), 1581–1590. 10.1194/jlr.D003715.

50. Mitroulis, I.; Kalafati, L.; Hajishengallis, G.; Chavakis, T. Myelopoiesis in the Context of Innate Immunity. J. Innate Immun. 2018, 10 (5–6), 365–372. 10.1159/000489406.

51. Qureshi, A.; Subathra, M.; Grey, A.; Schey, K.; Del Poeta, M.; Luberto, C. Role of Sphingomyelin Synthase in Controlling the Antimicrobial Activity of Neutrophils against Cryptococcus Neoformans. PLoS ONE 2010, 5 (12), e15587. 10.1371/journal.pone.0015587.

52. Hinkovska-Galcheva, V. Tz.; Boxer, L. A.; Mansfield, P. J.; Harsh, D.; Blackwood, A.; Shayman, J. A. The Formation of Ceramide-1-Phosphate during Neutrophil Phagocytosis and Its Role in Liposome Fusion. J. Biol. Chem. 1998, 273 (50), 33203–33209. 10.1074/jbc.273.50.33203.

53. Gutierrez, M. G.; Gonzalez, A. P.; Anes, E.; Griffiths, G. Role of Lipids in Killing Mycobacteria by Macrophages: Evidence for NF-κB-Dependent and -Independent Killing Induced by Different Lipids. Cell. Microbiol. 2009, 11 (3), 406–420. 10.1111/j.1462-5822.2008.01263.x.

54. Fadok, V. A.; Bratton, D. L.; Frasch, S. C.; Warner, M. L.; Henson, P. M. The Role of Phosphatidylserine in Recognition of Apoptotic Cells by Phagocytes. Cell Death Differ. 1998, 5 (7), 551–562. 10.1038/sj.cdd.4400404.

55. Brouwers, H.; Jónasdóttir, H. S.; Kuipers, M. E.; Kwekkeboom, J. C.; Auger, J. L.; Gonzalez-Torres, M.; López-Vicario, C.; Clària, J.; Freysdottir, J.; Hardardottir, I.; Garrido-Mesa, J.; Norling, L. V.; Perretti, M.; Huizinga, T. W. J.; Kloppenburg, M.; Toes, R. E. M.; Binstadt, B.; Giera, M.; Ioan-Facsinay, A. Anti-Inflammatory and Proresolving Effects of the Omega-6 Polyunsaturated Fatty Acid Adrenic Acid. J. Immunol. Baltim. Md 1950 2020, 205 (10), 2840–2849. 10.4049/jimmunol.1801653.

56. Norris, P. C.; Dennis, E. A. Omega-3 Fatty Acids Cause Dramatic Changes in TLR4 and Purinergic Eicosanoid Signaling. Proc. Natl. Acad. Sci. 2012, 109 (22), 8517–8522. 10.1073/pnas.1200189109.

57. Scott, A. J.; Post, J. M.; Lerner, R.; Ellis, S. R.; Lieberman, J.; Shirey, K. A.; Heeren, R. M. A.; Bindila, L.; Ernst, R. K. Host-Based Lipid Inflammation Drives Pathogenesis in Francisella Infection. Proc. Natl. Acad. Sci. 2017, 114 (47), 12596–12601. 10.1073/pnas.1712887114.

58. Kalinski, P. Regulation of Immune Responses by Prostaglandin E2. J. Immunol. 2012, 188 (1), 21–28. 10.4049/jimmunol.1101029.

59. Miek, L.; Jordan, P. M.; Günther, K.; Pace, S.; Beyer, T.; Kowalak, D.; Hoerr, V.; Löffler, B.; Tuchscherr, L.; Serhan, C. N.; Gerstmeier, J.; Werz, O. Staphylococcus Aureus Controls Eicosanoid and Specialized Pro-Resolving Mediator Production via Lipoteichoic Acid. Immunology 2022, 166 (1), 47–67. 10.1111/imm.13449.

60. Scher, J. U.; Pillinger, M. H. The Anti-Inflammatory Effects of Prostaglandins. J. Investig. Med. 2009, 57 (6), 703–708. 10.2310/JIM.0b013e31819aaa76.

61. Lebrero, P.; Astudillo, A. M.; Rubio, J. M.; Fernández-Caballero, L.; Kokotos, G.; Balboa, M. A.; Balsinde, J. Cellular Plasmalogen Content Does Not Influence Arachidonic Acid Levels or Distribution in Macrophages: A Role for Cytosolic Phospholipase A2γ in Phospholipid Remodeling. Cells 2019, 8 (8), 799. 10.3390/cells8080799.

62. Guerrini, V.; Gennaro, M. L. Foam Cells: One Size Doesn’t Fit All. Trends Immunol. 2019, 40 (12), 1163–1179. 10.1016/j.it.2019.10.002.

63. Olzmann, J. A.; Carvalho, P. Dynamics and Functions of Lipid Droplets. Nat. Rev. Mol. Cell Biol. 2019, 20 (3), 137–155. 10.1038/s41580-018-0085-z.

64. Guerrini, V.; Prideaux, B.; Khan, R.; Subbian, S.; Wang, Y.; Sadimin, E.; Pawar, S.; Ukey, R.; Singer, E. A.; Xue, C.; Gennaro, M. L. Heterogeneity of Foam Cell Biogenesis across Diseases. bioRxiv 2023, 2023.06.08.542766. 10.1101/2023.06.08.542766.

65. Agarwal, P.; Combes, T. W.; Shojaee-Moradie, F.; Fielding, B.; Gordon, S.; Mizrahi, V.; Martinez, F. O. Foam Cells Control Mycobacterium Tuberculosis Infection. Front. Microbiol. 2020, 11, 1394. 10.3389/fmicb.2020.01394.

66. Nicolaou, G.; Goodall, A. H.; Erridge, C. Diverse Bacteria Promote Macrophage Foam Cell Formation via Toll-like Receptor-Dependent Lipid Body Biosynthesis. J. Atheroscler. Thromb. 2012, 19 (2), 137–148. 10.5551/jat.10249.

67. D’Avila, H.; Melo, R. C. N.; Parreira, G. G.; Werneck-Barroso, E.; Castro-Faria-Neto, H. C.; Bozza, P. T. Mycobacterium Bovis Bacillus Calmette-Guérin Induces TLR2-Mediated Formation of Lipid Bodies: Intracellular Domains for Eicosanoid Synthesis In Vivo1. J. Immunol. 2006, 176 (5), 3087–3097. 10.4049/jimmunol.176.5.3087.

68. Denning, N.-L.; Aziz, M.; Gurien, S. D.; Wang, P. DAMPs and NETs in Sepsis. Front. Immunol. 2019, 10.

69. Gibson, M. S.; Domingues, N.; Vieira, O. V. Lipid and Non-Lipid Factors Affecting Macrophage Dysfunction and Inflammation in Atherosclerosis. Front. Physiol. 2018, 9.

70. Anderson, D. M. G.; Ablonczy, Z.; Koutalos, Y.; Hanneken, A. M.; Spraggins, J. M.; Calcutt, M. W.; Crouch, R. K.; Caprioli, R. M.; Schey, K. L. Bis(Monoacylglycero)Phosphate Lipids in the Retinal Pigment Epithelium Implicate Lysosomal/Endosomal Dysfunction in a Model of Stargardt Disease and Human Retinas. Sci. Rep. 2017, 7, 17352. 10.1038/s41598-017-17402-1.

71. Hullin-Matsuda, F.; Luquain-Costaz, C.; Bouvier, J.; Delton-Vandenbroucke, I. Bis(Monoacylglycero)Phosphate, a Peculiar Phospholipid to Control the Fate of Cholesterol: Implications in Pathology. Prostaglandins Leukot. Essent. Fatty Acids 2009, 81 (5), 313–324. 10.1016/j.plefa.2009.09.006.

72. Luquain-Costaz, C.; Lefai, E.; Arnal-Levron, M.; Markina, D.; Sakaï, S.; Euthine, V.; Makino, A.; Guichardant, M.; Yamashita, S.; Kobayashi, T.; Lagarde, M.; Moulin, P.; Delton-Vandenbroucke, I. Bis(Monoacylglycero)Phosphate Accumulation in Macrophages Induces Intracellular Cholesterol Redistribution, Attenuates Liver-X Receptor/ATP-Binding Cassette Transporter A1/ATP-Binding Cassette Transporter G1 Pathway, and Impairs Cholesterol Efflux. Arterioscler. Thromb. Vasc. Biol. 2013, 33 (8), 1803–1811. 10.1161/ATVBAHA.113.301857.

73. Bobryshev, Y. V.; Lord, R. S. A.; Golovanova, N. K.; Gracheva, E. V.; Zvezdina, N. D.; Prokazova, N. V. Phenotype Determination of Anti-GM3 Positive Cells in Atherosclerotic Lesions of the Human Aorta: Hypothetical Role of Ganglioside GM3 in Foam Cell Formation. Biochim. Biophys. Acta BBA - Mol. Basis Dis. 2001, 1535 (2), 87–99. 10.1016/S0925-4439(00)00076-4.

74. Prokazova, N. V.; Mikhailenko, I. A.; Bergelson, L. D. Ganglioside GM3 Stimulates the Uptake and Processing of Low Density Lipoproteins by Macrophages. Biochem. Biophys. Res. Commun. 1991, 177 (1), 582–587. 10.1016/0006-291X(91)92023-D.

75. Glaros, E. N.; Kim, W. S.; Quinn, C. M.; Wong, J.; Gelissen, I.; Jessup, W.; Garner, B. Glycosphingolipid Accumulation Inhibits Cholesterol Efflux via the ABCA1/Apolipoprotein A-I Pathway. J. Biol. Chem. 2005, 280 (26), 24515–24523. 10.1074/jbc.M413862200.

76. Akgoc, Z.; Sena-Esteves, M.; Martin, D. R.; Han, X.; d’Azzo, A.; Seyfried, T. N. Bis(Monoacylglycero)Phosphate: A Secondary Storage Lipid in the Gangliosidoses. J. Lipid Res. 2015, 56 (5), 1005–1006. 10.1194/jlr.M057851.

77. Brok, M. H. den; Raaijmakers, T. K.; Collado-Camps, E.; Adema, G. J. Lipid Droplets as Immune Modulators in Myeloid Cells. Trends Immunol. 2018, 39 (5), 380–392. 10.1016/j.it.2018.01.012.

78. Almeida, P. E.; Silva, A. R.; Maya-Monteiro, C. M.; Töröcsik, D.; D′Ávila, H.; Dezsö, B.; Magalhães, K. G.; Castro-Faria-Neto, H. C.; Nagy, L.; Bozza, P. T. Mycobacterium Bovis Bacillus Calmette-Guérin Infection Induces TLR2-Dependent Peroxisome Proliferator-Activated Receptor γ Expression and Activation: Functions in Inflammation, Lipid Metabolism, and Pathogenesis1. J. Immunol. 2009, 183 (2), 1337–1345. 10.4049/jimmunol.0900365.

79. Stewart, C. R.; Stuart, L. M.; Wilkinson, K.; van Gils, J. M.; Deng, J.; Halle, A.; Rayner, K. J.; Boyer, L.; Zhong, R.; Frazier, W. A.; Lacy-Hulbert, A.; Khoury, J. E.; Golenbock, D. T.; Moore, K. J. CD36 Ligands Promote Sterile Inflammation through Assembly of a Toll-like Receptor 4 and 6 Heterodimer. Nat. Immunol. 2010, 11 (2), 155–161. 10.1038/ni.1836.

80. Jongstra-Bilen, J.; Zhang, C. X.; Wisnicki, T.; Li, M. K.; White-Alfred, S.; Ilaalagan, R.; Ferri, D. M.; Deonarain, A.; Wan, M. H.; Hyduk, S. J.; Cummins, C. L.; Cybulsky, M. I. Oxidized Low-Density Lipoprotein Loading of Macrophages Downregulates TLR-Induced Proinflammatory Responses in a Gene-Specific and Temporal Manner through Transcriptional Control. J. Immunol. 2017, 199 (6), 2149–2157. 10.4049/jimmunol.1601363.

81. Djambazova, K. V.; Gibson-Corley, K. N.; Freiberg, J. A.; Caprioli, R. M.; Skaar, E. P.; Spraggins, J. M. MALDI TIMS IMS Reveals Ganglioside Molecular Diversity within Murine S. Aureus Soft Tissue Abscesses. ChemRxiv November 21, 2023. 10.26434/chemrxiv-2023-whtvz.

82. Carrillo-Marquez, M. A.; Hulten, K. G.; Hammerman, W.; Mason, E. O.; Kaplan, S. L. USA300 Is the Predominant Genotype Causing Staphylococcus Aureus Septic Arthritis in Children. Pediatr. Infect. Dis. J. 2009, 28 (12), 1076. 10.1097/INF.0b013e3181adbcfe.

83. Kawamoto, T.; Kawamoto, K. Preparation of Thin Frozen Sections from Nonfixed and Undecalcified Hard Tissues Using Kawamot’s Film Method (2012). In Skeletal Development and Repair: Methods and Protocols; Hilton, M. J., Ed.; Methods in Molecular Biology; Humana Press: Totowa, NJ, 2014; pp 149–164. 10.1007/978-1-62703-989-5_11.

84. Spraggins, J. M.; Djambazova, K. V.; Rivera, E. S.; Migas, L. G.; Neumann, E. K.; Fuetterer, A.; Suetering, J.; Goedecke, N.; Ly, A.; Van de Plas, R.; Caprioli, R. M. High-Performance Molecular Imaging with MALDI Trapped Ion-Mobility Time-of-Flight (timsTOF) Mass Spectrometry. Anal. Chem. 2019, 91 (22), 14552–14560. 10.1021/acs.analchem.9b03612.

85. Kawamoto, K.; Suzuki, T.; Nagano, T.; Kawamoto, T.; Gomi, K. A Study of Bone Formation around Titanium Implants Using Frozen Sections. J. Hard Tissue Biol. 2021, 30 (2), 165–174. 10.2485/jhtb.30.165.

86. Colley, M. E.; Allen, J.; Djambazova, K. V.; Kruse, A.; melissa.a.farrow; Spraggins, J. Bulk Untargeted LC-MS/MS Lipidomics. 2023.

87. Matyash, V.; Liebisch, G.; Kurzchalia, T. V.; Shevchenko, A.; Schwudke, D. Lipid Extraction by Methyl-Tert-Butyl Ether for High-Throughput Lipidomics. J. Lipid Res. 2008, 49 (5), 1137–1146. 10.1194/jlr.D700041-JLR200.

88. Vasilopoulou, C. G.; Sulek, K.; Brunner, A.-D.; Meitei, N. S.; Schweiger-Hufnagel, U.; Meyer, S. W.; Barsch, A.; Mann, M.; Meier, F. Trapped Ion Mobility Spectrometry and PASEF Enable In-Depth Lipidomics from Minimal Sample Amounts. Nat. Commun. 2020, 11 (1), 331. 10.1038/s41467-019-14044-x.

89. Tsugawa, H.; Ikeda, K.; Takahashi, M.; Satoh, A.; Mori, Y.; Uchino, H.; Okahashi, N.; Yamada, Y.; Tada, I.; Bonini, P.; Higashi, Y.; Okazaki, Y.; Zhou, Z.; Zhu, Z.-J.; Koelmel, J.; Cajka, T.; Fiehn, O.; Saito, K.; Arita, M.; Arita, M. A Lipidome Atlas in MS-DIAL 4. Nat. Biotechnol. 2020, 38 (10), 1159–1163. 10.1038/s41587-020-0531-2.

90. Sud, M.; Fahy, E.; Cotter, D.; Brown, A.; Dennis, E. A.; Glass, C. K.; Merrill, A. H.; Murphy, R. C.; Raetz, C. R. H.; Russell, D. W.; Subramaniam, S. LMSD: LIPID MAPS Structure Database. Nucleic Acids Res. 2007, 35 (Database issue), D527–D532. 10.1093/nar/gkl838.

91. Liebisch, G.; Fahy, E.; Aoki, J.; Dennis, E. A.; Durand, T.; Ejsing, C. S.; Fedorova, M.; Feussner, I.; Griffiths, W. J.; Köfeler, H.; Merrill, A. H.; Murphy, R. C.; O’Donnell, V. B.; Oskolkova, O.; Subramaniam, S.; Wakelam, M. J. O.; Spener, F. Update on LIPID MAPS Classification, Nomenclature, and Shorthand Notation for MS-Derived Lipid Structures. J. Lipid Res. 2020, 61 (12), 1539–1555. 10.1194/jlr.S120001025.

92. Ulmer, C. Z.; Koelmel, J. P.; Ragland, J. M.; Garrett, T. J.; Bowden, J. A. LipidPioneer: A Comprehensive User-Generated Exact Mass Template for Lipidomics. J. Am. Soc. Mass Spectrom. 2017, 28 (3), 562–565. 10.1007/s13361-016-1579-6.

