## Supplementary material for "*In situ* lipidomics of *Staphylococcus aureus* osteomyelitis using imaging mass spectrometry": Figure S1

### Table of Contents

Figures

S1) All section replicates for pre-IMS fluorescence microscopy and post-IMS histology.

S2) All section replicates for *k*-means clustering analysis and alterations to clustering results following *m/z* feature loss and tissue wash.

S3) Extracted average mass spectra for select *k*-means clusters.

S4) Ion images comparing unwashed and salt-washed data.

S5) Additional clustering results and lipid intensity data for the analysis of single abscesses.

S6) Intensity comparison of lipids between infected and mock-infected bone marrow.

S7) Validation of the identification of bis(monoacylglycero)phosphates in the abscess border using ion mobility.

Tables **(In Excel File)**

S1) Table of accurate mass measurements, molecular identifications, and supplemental LC-MS/MS information for all lipids referenced in this study (Positive Ion Mode).

S2) Table of accurate mass measurements, molecular identifications, and supplemental LC-MS/MS information for all lipids referenced in this study (Negative Ion Mode).

S3) Table of lipid markers of positive ion clusters largely affected by infection.

S4) Table of lipid markers of positive ion clusters largely unaffected by infection.

S5) Table of lipid markers of negative ion clusters largely affected by infection.

S6) Table of lipid markers of negative ion clusters largely unaffected by infection.

S7) Table of detected lipids in staphylococcal abscess communities.

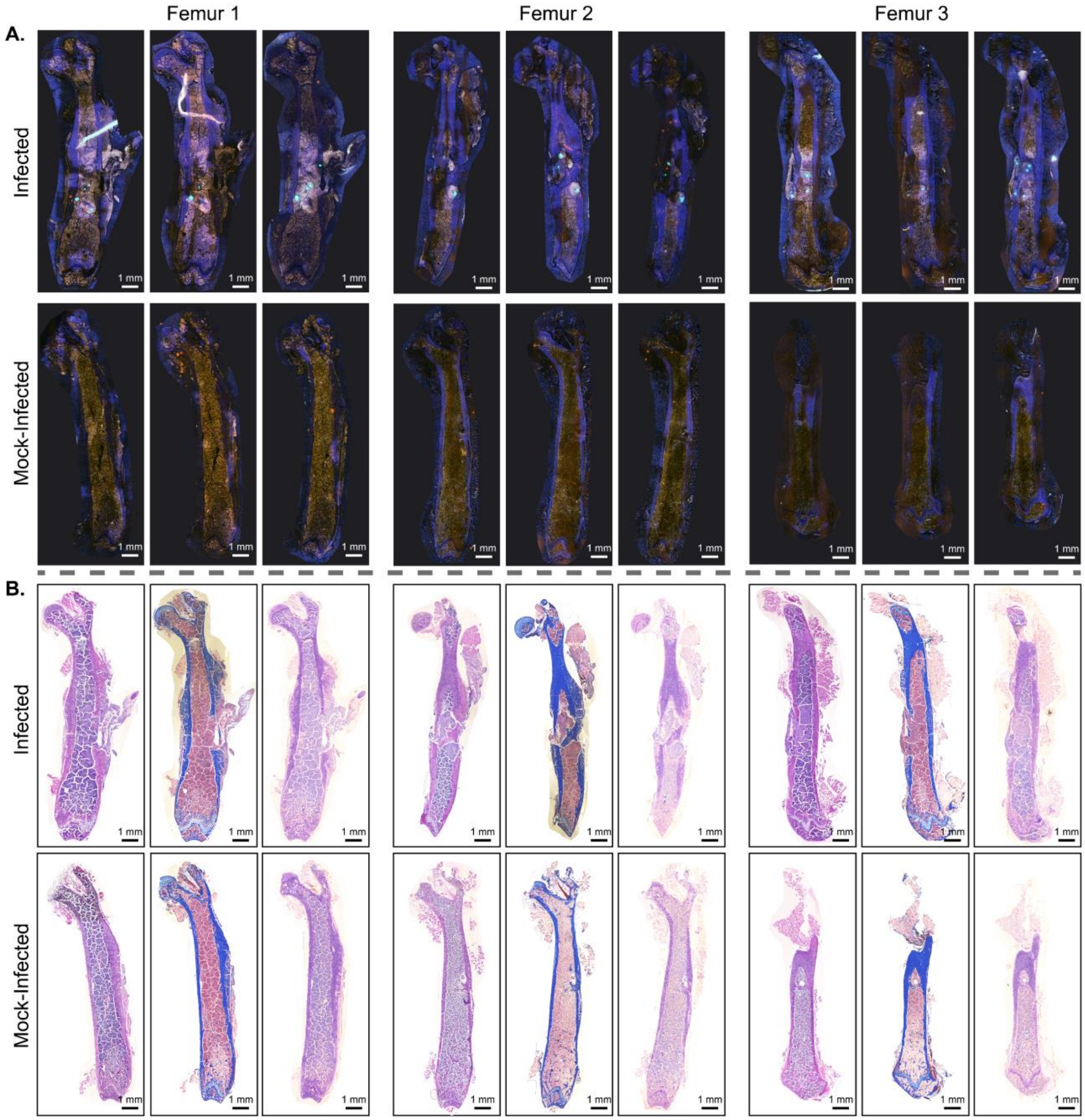


**Figure S1. Pre-IMS fluorescence images and post-IMS histological stains for all infected and mock-infected section replicates in the main dataset are shown.** (A) Fluorescence images indicate that SACs are present in each infected femur and also show a more intense autofluorescence signal in infected bone marrow compared to mock-infected bone marrow. (B) Hematoxylin and eosin (H&E, Left), Masson's Trichrome (MTC, Middle), and Oil Red O (ORO, Right) stains are leveraged to identify large tissue features like surrounding soft tissue, calcified bone, and fat deposits. ORO images are saturated more to reveal the subtle orange-red staining. ORO staining is less pronounced than expected in the surrounding adipose tissue and the fibrotic abscess border, and the MALDI IMS experiment may have been responsible.

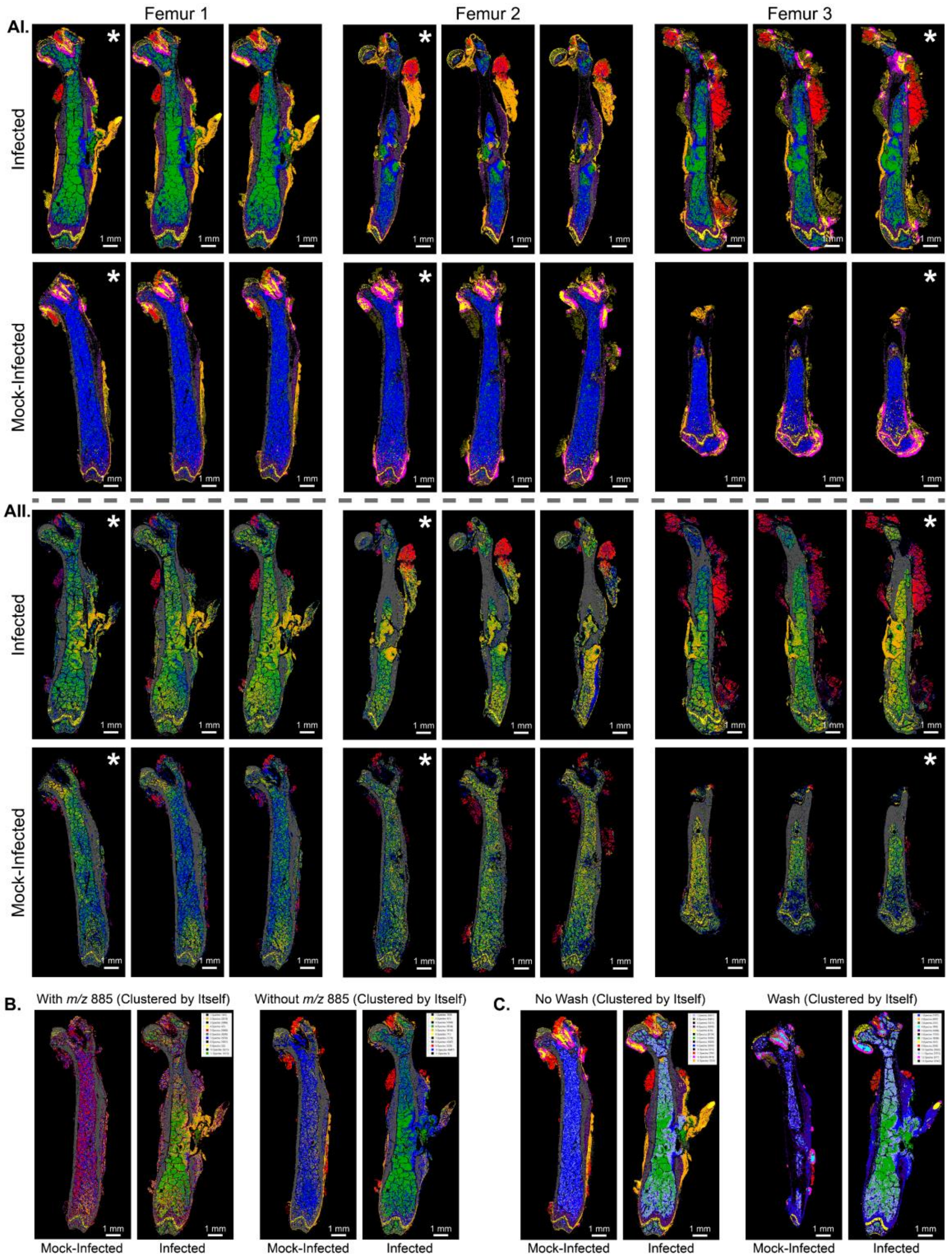


**Figure S2. Clustering results for all section replicates from the main *k*-means analysis and results from additional clustering experiments are reported.** (A) *k*-Means clustering for all (AI) positive and (AII) negative ion data reveal the same clustering patterns as described in **Figure 3** (* denotes replicates shown in main figure). (B,C) When an infected and mock-infected pair are clustered separately from the entire replicate dataset ("Clustered by Itself"), the clustering results change due to less biological and tissue variation from all samples. (B) When the most abundant negative ion, *m/z* 885.550, is removed from the clustering analysis, the negative clusters become more defined and more similar to positive ion analysis. (C) When salt adducts are removed from the tissue, the clustering is similar to its no washed counterpart. The main difference seems to be the combination of the outside cluster (previously orange) with the bone marrow (blue) found in infected and mock-infected tissue, most likely a result of the loss of sodium on the outside of the femur.

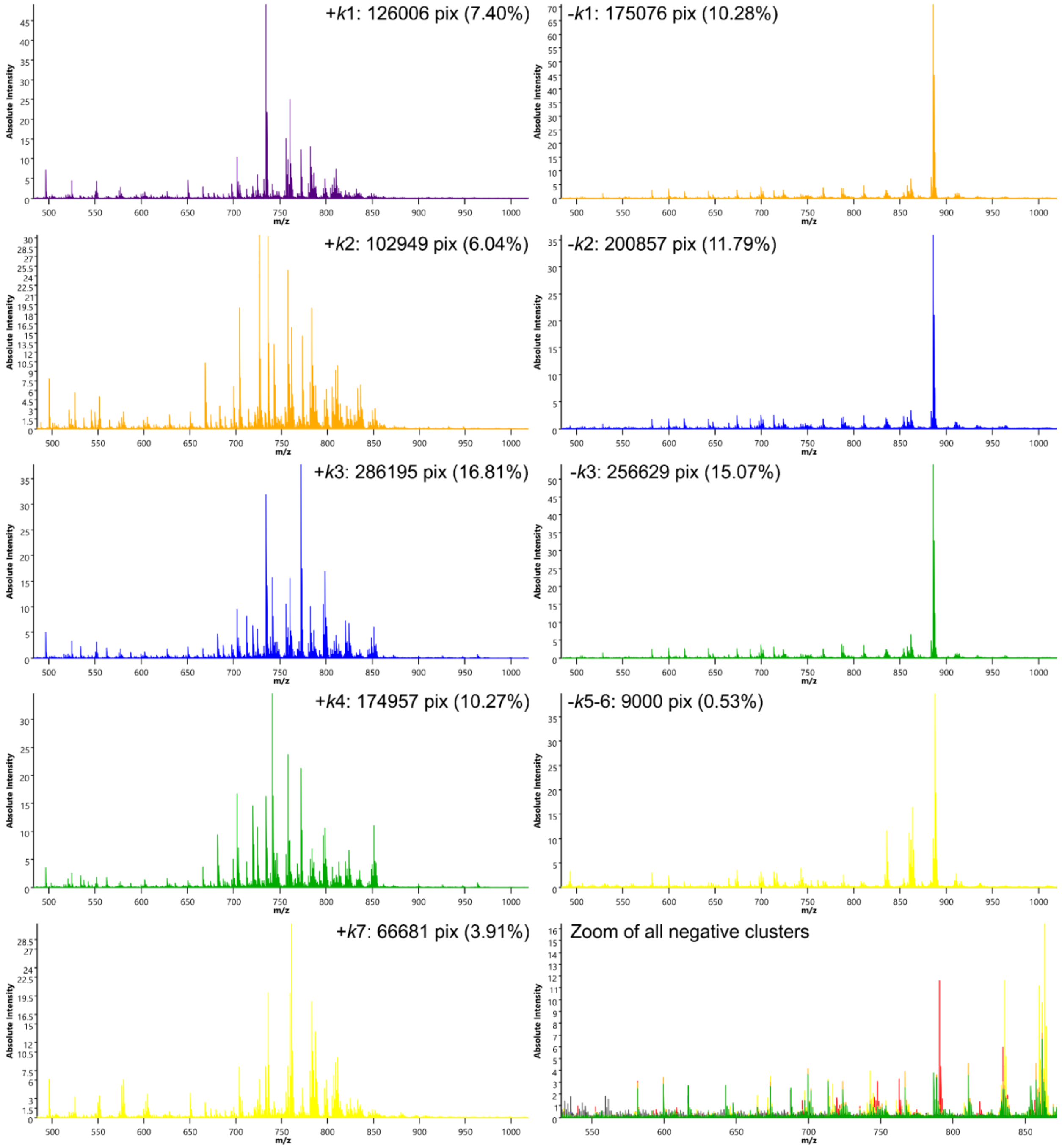


**Figure S3. Extracted average mass spectra for select clusters from positive ion (Left) and negative ion (Right) analysis highlight the molecular heterogeneity detected in clusters correlating to bone- and infection-associated tissue types.** The total number of pixels across all infected and mock-infected samples that are included in the average spectrum of each cluster are reported. The percentage of pixels in a cluster from the total imaged pixels are also reported to indicate a relative quantity of the corresponding tissue types throughout the samples.

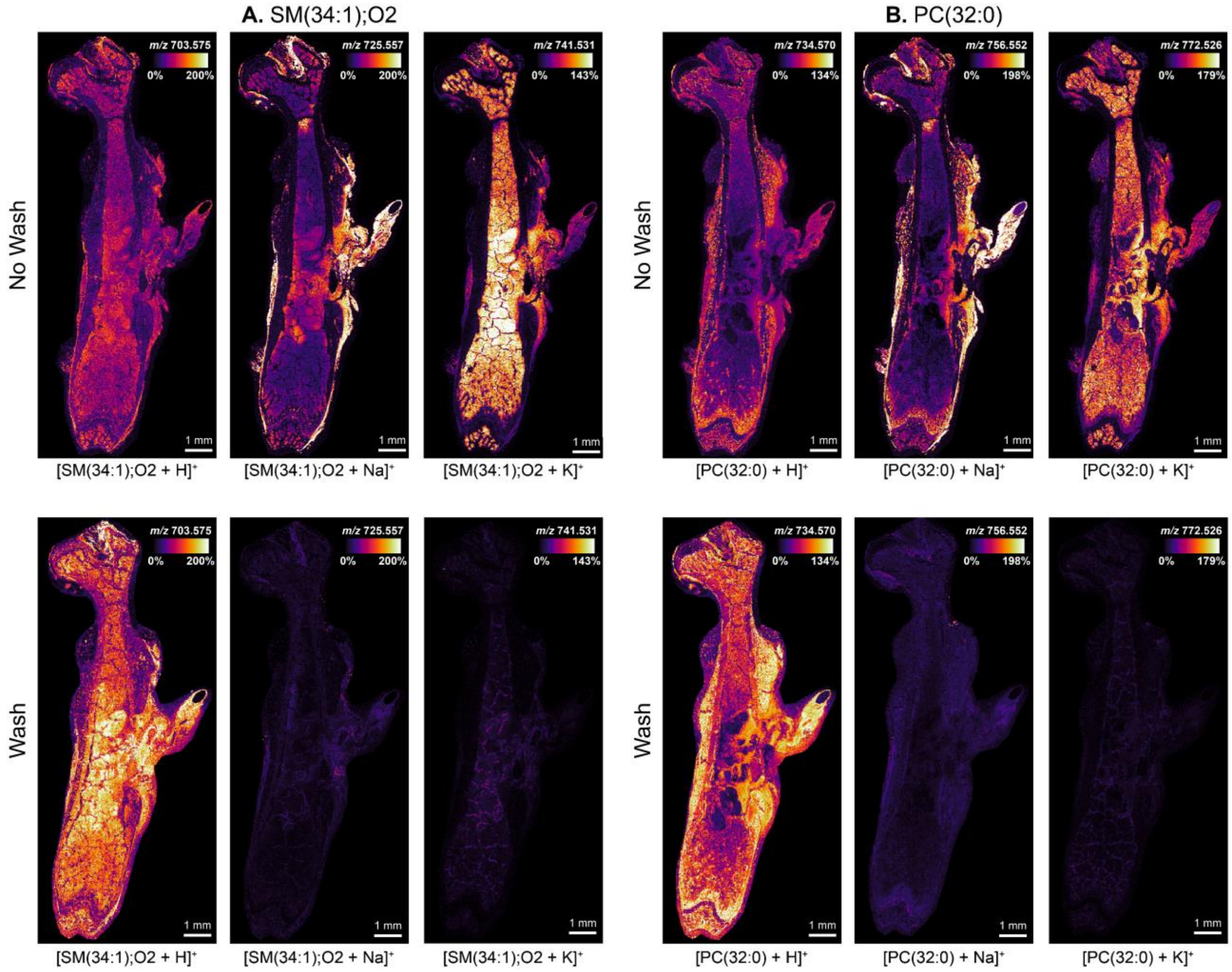


**Figure S4. Ion images reveal how salt-washing reduces adduct competition and simplifies distributions.** The adducts of the most abundant (A) sphingolipid and (B) glycerophospholipid are compared between unwashed and salt-washed samples. Sodiated ions are largely abundant on the outside of the intramedullary cavity, and potassiated products are largely abundant on the inside (Top), which recapitulates the trends discovered in the lipid marker analysis (**Figure 3 and Table S3**). All salt adducts are virtually depleted, and protonation yields the truest distribution of the lipids (Bottom), which appears to be the sum of all unwashed adducts. Ion images of the same adducts or *m/z* are on the same hotspot intensity scale to compare abundances between the unwashed and salt-washed data. This salt wash experiment is also fundamental for confirming identifications of different adduct species.

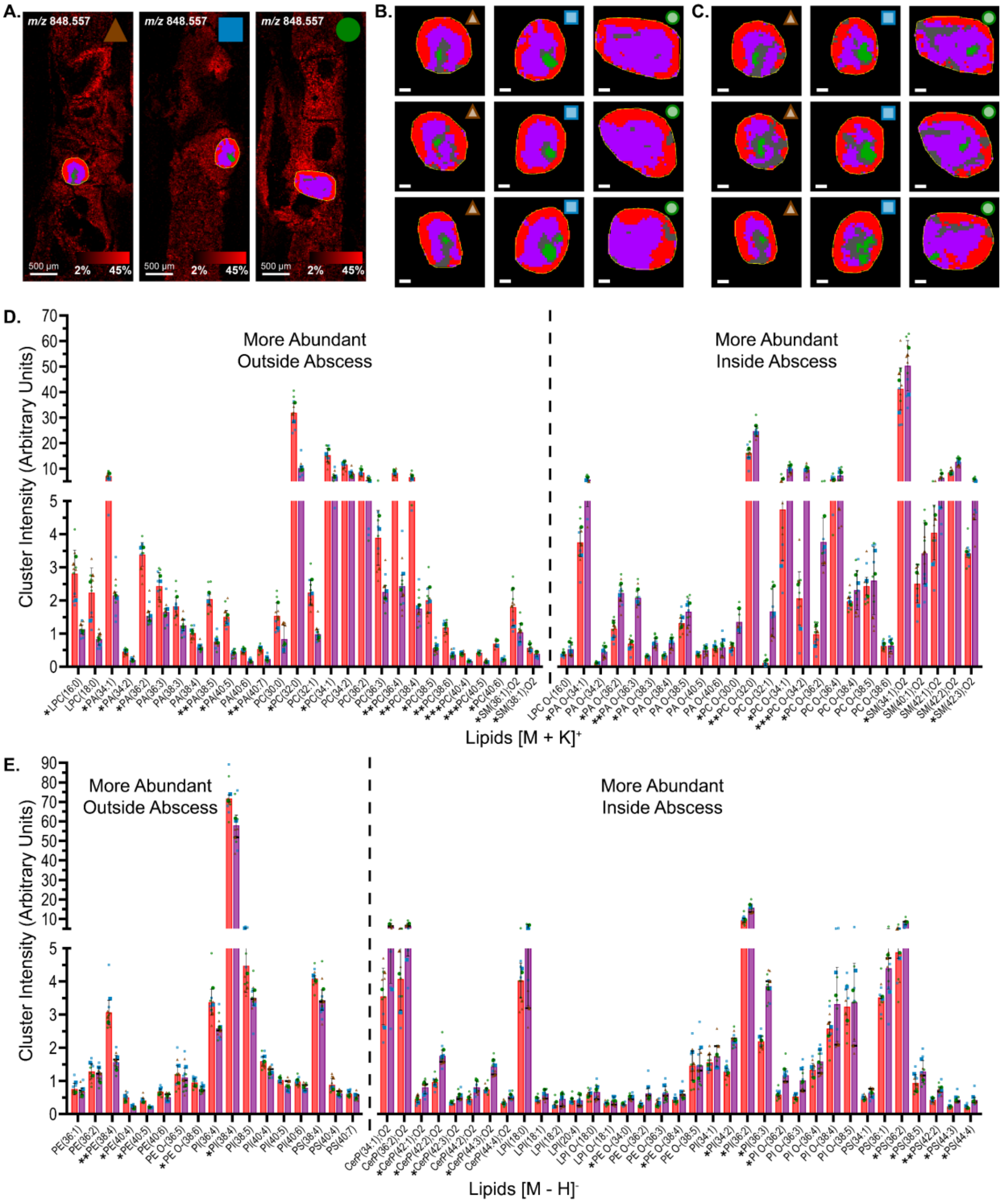


**Figure S5. Section replicate clustering results and insignificant lipid intensity changes between abscess regions are reported.** (A) The location of the selected abscess within each intramedullary cavity is shown. (B,C) All nine clustering replicates are reported for each (B) positive and (C) negative ion analysis. Scale bars represent 100 µm. (D,E) Bar graphs show statistically significant and insignificant differences in lipid abundances between the outside abscess pixels and inside abscess pixels for each (D) positive and (E) negative ion analysis. Similar to **Figure 5**, the mean, with biological standard deviation, for three femurs is displayed as a bar with colors corresponding to the derived cluster. Section replicate measurements and replicate averages (one femur) are displayed on the graph. A paired T-test is used to test statistical differences, with *<0.05, **<0.01, ***<0.005 denoted below the graph on the x-axis labels. Here, additional insignificant lipids are reported and follow similar trends but are not biologically significant due to large abscess and replicate variation. Lipids are also sorted in alphabetical order rather than mean intensity.

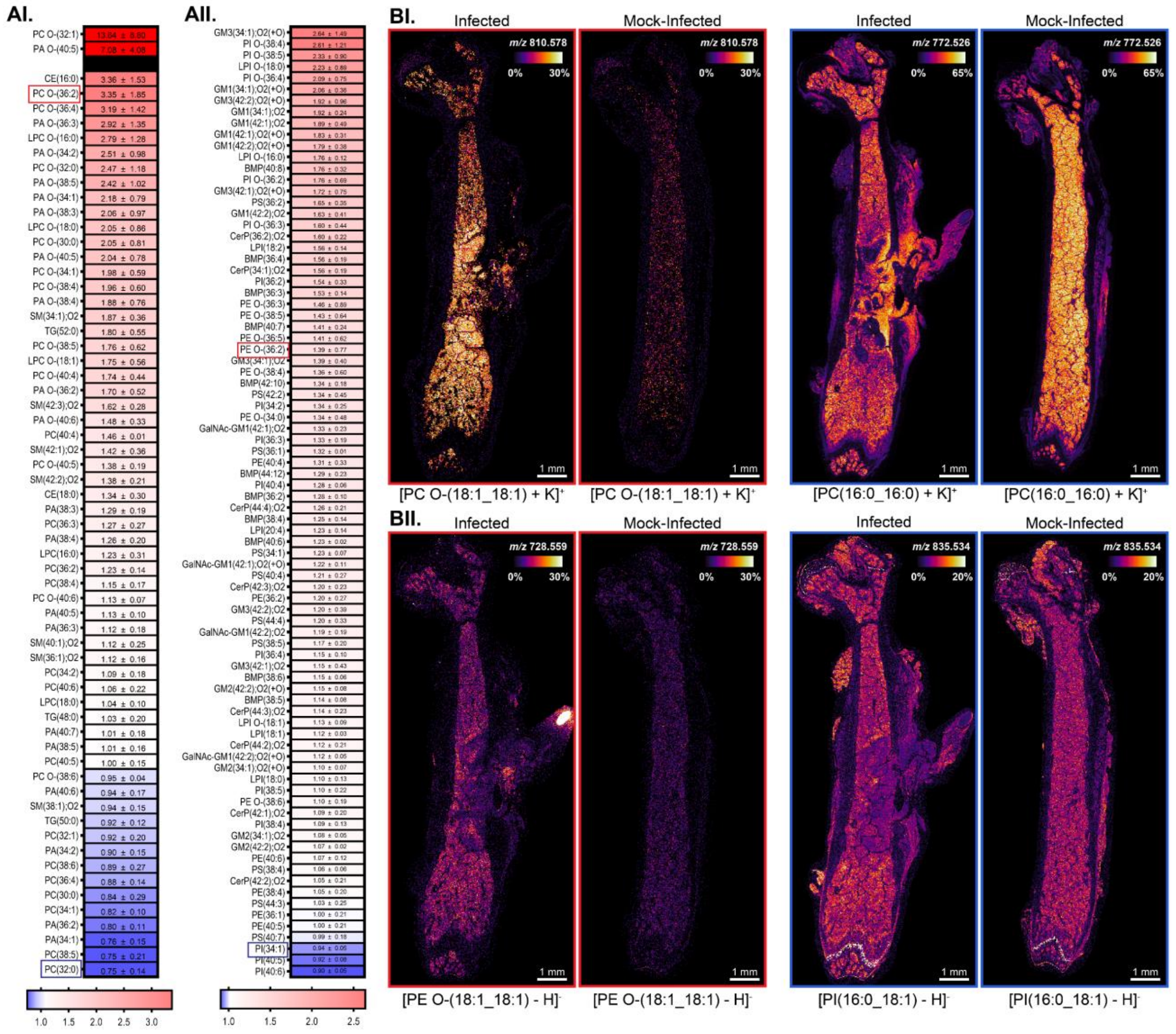


**Figure S6. Intensity fold changes and representative ion images indicate how the abundances of lipids globally change between infected and mock-infected bone marrow.** (A) Lipids are sorted based on their intensity fold change from the mock-infected bone marrow for (AI) positive ion (+K) and (AII) negative ion (-H) data. For example, PC O-(36:2) in infected bone marrow is 3.35x more intense than mock-infected. Two lipids in positive ion mode, PC O-(32:1) and PA O-(40:5) are outliers from the scaling. Means and biological standard deviations are reported on the color scale. (B) Representative ion images are shown for (BI) positive and (BII) negative ion analysis.

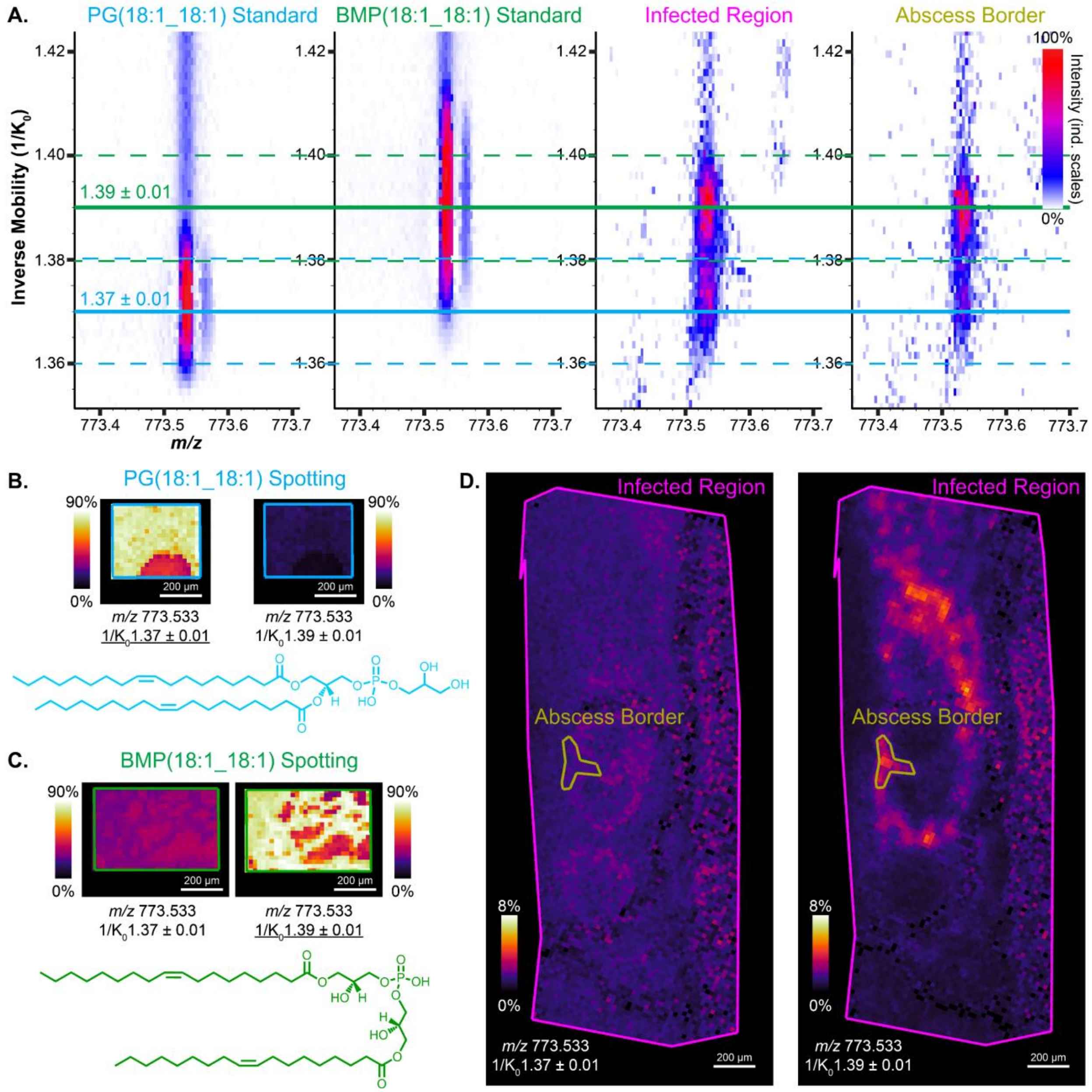


**Figure S7. MALDI TIMS imaging confirms identification of BMPs in the abscess border.** (A) Heat maps are extracted from imaging data of two lipid standards and two regions of an infected femur, and they show differences in inverse mobility distributions of *m/z* 773.533. The PG(18:1_18:1) standard and BMP(18:1_18:1) standard separate by TIMS, and their respective $1/K_0$ values can be tracked in the infection data. (B, C, and D) Ion images of selected regions are displayed to recapitulate heat map results. (B) The blue box is where the PG standard is spotted, and *m/z* is most intense at a $1/K_0$ of 1.37. (C) The green box is where the BMP standard is spotted, and *m/z* is most intense at a $1/K_0$ of 1.39. There is some signal in the other mobility window because the peaks are not entirely baseline resolved. (D) Most of the signal in the abscess border is coming from the ion with a $1/K_0$ of 1.39, which would be endogenous BMP(36:2). Endogenous PG(36:2) could be present in other regions of the tissue, but the intensity is too low to distinguish from noise. All data are normalized together, and standards are more concentrated than the endogenous lipids which explains the differences in ion image scaling.
